# Viral Capsid-Membrane Interactions Propel Non-Brownian Movements of Non-enveloped Reoviruses during Entry

**DOI:** 10.64898/2026.01.21.700940

**Authors:** Mengchi Jiao, Gregory R. Cantrall, Yanqi Yu, Anthony J. Snyder, Steven M. Abel, Pranav Danthi, Yan Yu

## Abstract

Understanding how non-enveloped viruses breach host cell membranes is critical for developing strategies to block viral entry, a key step in infection. Despite extensive study, how viral capsids and host lipid membranes dynamically cooperate during membrane penetration remains poorly defined. Here, using reovirus as a model non-enveloped virus and planar model membranes, we identify previously unrecognized non-Brownian membrane motions of infectious subvirion particles (ISVPs) by single-virus tracking. We then integrate experiments with computational modeling to dissect the stepwise, processive capsid–membrane interactions encoded in these distinct dynamics. We show that ISVP motion transitions from an initial phase of directed translocation to progressively confined diffusion. This behavior reflects a multistep entry mechanism in which initial capsid–membrane contact triggers release of the membrane-active μ1N peptide. As μ1N accumulates within the bilayer, it generates membrane-associated viral retention sites that promote further virus adsorption and increasingly constrain particle mobility. By directly visualizing these motion signatures, we resolve transient and cooperative capsid–membrane interactions that are difficult to capture using conventional biochemical approaches. Together, these findings provide new insight into early membrane penetration events of non-enveloped viruses.

**SIGNIFICANCE STATEMENT:** Non-enveloped viruses must penetrate host cell membranes to initiate infection, yet the dynamic mechanisms by which viral capsids engage and remodel lipid membranes remain poorly understood. By directly visualizing the motion of individual reovirus particles on model membranes, we uncover previously unrecognized non-Brownian dynamics that encode stepwise capsid–membrane interactions during viral entry. Integrating single-virus tracking with computational modeling reveals how peptide-mediated membrane remodeling feeds back to regulate viral engagement. This work provides the first direct visualization of processive, stepwise capsid–membrane interactions during non-enveloped virus entry and establishes viral particle dynamics as a quantitative readout for dissecting membrane penetration mechanisms.

## INTRODUCTION

Non-enveloped viruses, which lack a lipid membrane surrounding the virus particle, face the challenge of overcoming the host membrane barrier to enter cells. Unlike enveloped viruses that deliver their genomic cargo by fusing their viral envelope with the host membrane, non-enveloped viruses rely on alternate mechanisms to initiate infection. Despite their phylogenetic diversity, studies suggest a conserved strategy among non-enveloped viruses for breaching the host membrane (*1, 2*). In most cases, conformational changes in the viral capsid facilitate interactions between capsid components and the host membrane. These interactions often result in membrane permeabilization and subsequent delivery of the viral genome to its target cellular compartment for infection.

Mammalian orthoreovirus (reovirus) serves as a powerful model to investigate this process. Reovirus is a prototypical non-enveloped virus from the *Spinareoviridae* family within the *Reovirales* order, which includes clinically significant pathogens such as rotavirus, Coltivirus, and Bluetongue virus (BTV). Reovirus particles consist of two concentric protein shells, with the inner shell encasing a double-stranded RNA genome (*3*). During infection, the outer capsid undergoes partial disassembly, forming infectious subvirion particles (ISVPs) (Fig. 1A). ISVPs subsequently convert to a conformationally altered particle, the ISVP*, and release membrane-active amphiphilic peptides, which insert into host membranes, likely creating perforations that enable the delivery of viral capsids into the cytoplasm, where they complete the remainder of the infectious cycle (*4–6*).

**Figure 1.**
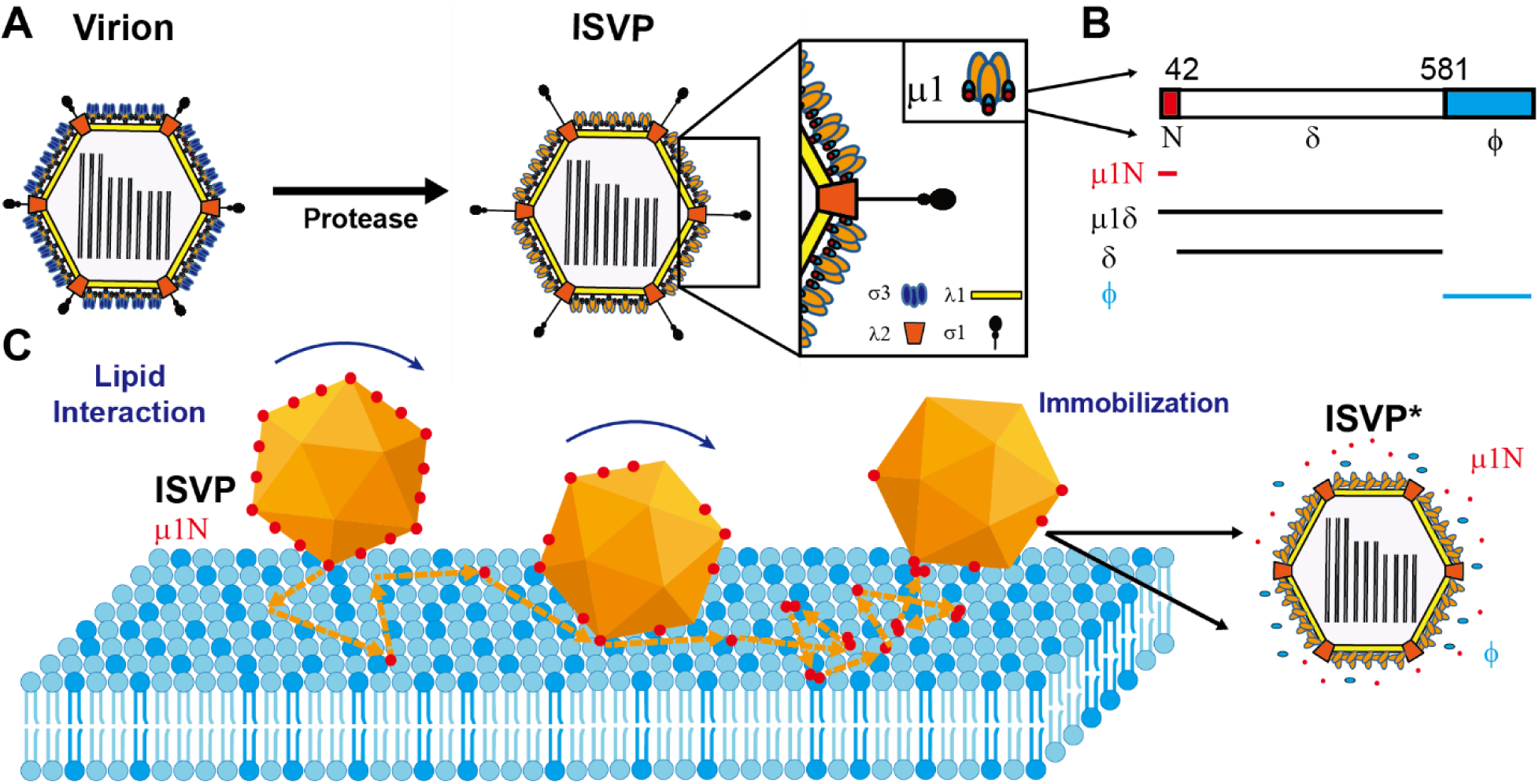
Schematic illustration of reovirus infectious subvirion particles (ISVPs) interaction with the planar-supported lipid bilayer. (A) Schematic illustration of the transition from Reovirus virion to ISVP and ISVP*. (B) Schematic illustration of cleavage fragments of the μ1 protein formed during ISVP lipid interactions. (C) Schematic illustration of peptide release from ISVP and embedded into the supported lipid bilayer during ISVP lipid interaction. At the same, the ISVP transitioned into ISVP* and immobilized on the membrane.

Previous work, including ours, has demonstrated that interactions between reovirus ISVPs and lipid membranes are critical for peptide release and subsequent membrane disruption (*7–9*). Using giant unilamellar vesicles (GUVs) as a model system, we recently showed that reovirus ISVPs exhibit dynamic interactions with the membrane, inducing local lipid deformations and capsid aggregation (*10*). These observations suggest a highly coordinated sequence of binding and capsid uncoating events. However, the specific roles of viral peptides, capsid dynamics, and lipid membranes in driving these interactions remain poorly understood.

Single-particle tracking offers a direct and widely used method to elucidate viral dynamics at the host membrane (*11, 12*). While this technique has been applied extensively to investigate the entry processes of enveloped viruses (*13–22*), it has surprisingly not been used to study the membrane interactions of reovirus or other non-enveloped viruses.

In this study, we applied a single-virus quantitative imaging approach with planar-supported lipid bilayers, combined with computational modeling, to reveal the complex interactions that govern reovirus–lipid membrane engagement. Our analysis resolved a stepwise sequence governing reovirus–membrane interactions. After contacting the bilayer, ISVPs exhibit coupled translational and rotational motion along the surface. Concurrently, viral peptides are released and inserted into the bilayer, where they accumulate locally to form transient retention sites that increase the probability of capturing virus particles. Over time, trajectories shift from diffusive to increasingly confined, which likely results in membrane breach required for entry. This study provides the first visualization of the stepwise interactions between reovirus particles, viral peptides, and lipid membranes during the membrane disruption process. Our findings offer critical insights into the entry mechanisms of non-enveloped viruses and lay the groundwork for comparative studies on how other non-enveloped viruses interact with host membranes during cell entry.

## MATERIALS and METHODS

### Materials and reagents

Joklik’s minimal essential medium was purchased from Lonza (Walkersville, MD). Fetal bovine serum (FBS) was purchased from Life Technologies (Carlsbad, CA). L-glutamine, Penicillin, and streptomycin were purchased from Invitrogen (Carlsbad, CA). Vertrel-XF specialty fluid was purchased from Dupont (Wilmington, DE). N-p-tosyl-L-lysine chloromethyl ketone (TLCK)-treated chymotrypsin were purchased from (Worthington Biochemical, Lakewood, NJ). CF®568 succinimidyl ester (CF568-NHS) was purchased from Biotium (Fremont, CA). 0.1 µm TetraSpeck^TM^ fluorescent microspheres and Qdot™ 705 streptavidin conjugate were purchased from ThermoFisher Scientific (Waltham, MA). 4-(2-hydroxyethyl)-1-piperazineethanesulfonic acid (HEPES), albumin from bovine serum (BSA), amphotericin B, phenylmethylsulfonyl fluoride (PMSF), and Amicon Ultra filters (30K) were purchased from MilliporeSigma (St. Louis, MO). 1,2-dioleoyl-sn-glycero-3-phosphocholine (DOPC), 1,2-dioleoyl-sn-glycero-3-phosphoethanolamine (DOPE), 1,2-dioleoyl-sn-glycero-3-phosphoethanolamine-N-(lissamine rhodamine B sulfonyl) (RhB-DOPE), 1,2-dioleoyl-sn-glycero-3-phosphoethanolamine-N-[(dipyrrometheneboron difluoride)butanoyl] (TopFluor®-PE (Bodipy-PE)), and the mini-extruder were purchased from Avanti Polar Lipids, Inc (Alabaster, AL). Ultrapure water (18.2 MΩ·cm) was used in all experiments. Dialysis buffer (10 mM Tris-HCl, 15 mM MgCl_2_, and 150 mM NaCl, pH 7.4), HEPES buffer (2 mM HEPES, 2 mM NaCl, pH 7.4), and sodium bicarbonate buffer (50 mM NaHCO_3_, pH 8.5) were prepared in the lab.

### Cells and viruses

Spinner-adapted murine L929 (L) cells were grown at 37°C in Joklik’s minimal essential medium supplemented with 5% FBS, 2 mM L-glutamine, 100 U/mL penicillin, 100 μg/mL streptomycin, and 25 ng/mL amphotericin B. Reovirus type 1 Lang (T1L), T1L containing the type 3 Dearing M2 gene segment (T1L/T3DM2), and T1L/T3DM2 D371A were generated by plasmid-based reverse genetics (*23–26*).

### Virus purification

Viruses were propagated and purified as previously described (*25, 27*). Briefly, L cells infected with second- or third-passage reovirus stocks were lysed by sonication. Virus particles were extracted from lysates using Vertrel-XF specialty fluid (*28*). The extracted particles were layered onto 1.2 to 1.4 g/mL CsCl step gradients. The gradients were then centrifuged at 187,000 × g for 4 h at 4°C in a SW 41 Ti rotor (Beckman Coulter, Brea, CA). Bands corresponding to purified virus particles (∼1.36 g/mL) were isolated and dialyzed into the dialysis buffer (*29*). Following dialysis, the particle concentration was determined by measuring the optical density of the purified virus stocks at 260 nm (OD260; 1 unit at OD260 = 2.1 ×10^12^ particles/mL) (*29*).

### Conjugation of reovirus with fluorescent dyes and biotin

Purified reovirus was labeled using CF568-NHS dye and biotin-NHS. Briefly, virions (1 ×10^13^ particles/mL in PBS) were diluted into fresh sodium bicarbonate buffer and incubated with 10 μM CF568-NHS and biotin-NHS (1 μM) for 90 min at room temperature in the dark. After 90 min, the reaction was quenched by the addition of dialysis buffer, and the labeled virions were layered onto 1.2- to 1.4 g/mL CsCl step gradients and repurified as described above.

### Generation of infectious subvirion particles (ISVPs)

T1L or T1L/T3DM2 virions (2 ×10^12^ particles/mL) were digested with 200 μg/mL TLCK-treated chymotrypsin in a total volume of 100 μL for 1 h at 37°C. After 1 h, the reaction mixtures were incubated on ice for 20 min and quenched by the addition of 1 mM PMSF. The generation of ISVPs was confirmed by SDS-PAGE and Coomassie staining (*8*).

### Generation of ISVP* supernatant

The supernatant of pre-converted ISVP*s was generated as previously described (*5, 30*). Briefly, T1L ISVPs (2 ×10^12^ particles/mL) were incubated at 52°C for 5 min. The heat-inactivated virus (no spin) was then centrifuged at 16,000 × g for 10 min at 4°C to pellet particles. The supernatant was assumed to contain released peptides μ1N along with Φ.

### Fluorescence microscopy

Epifluorescence images and total internal reflection fluorescence (TIRF) microscopy images were acquired using a Nikon Eclipse Ti-E inverted microscope equipped with a Nikon 100×/1.49 N.A. Oil-immersion TIRF objective and a Hamamatsu ORCA-Fusion digital CMOS camera. Glass coverslips for imaging were cleaned by sonication with 70% ethanol aqueous solution and rinsed with Milli-Q^®^ water before being assembled into the imaging chamber.

### Planar-supported lipid bilayer experiments

#### (a) Large unilamellar vesicles (LUVs) formation

LUVs of 100 nm in diameter were made using the extrusion method (*31, 32*). To prepare the LUVs, 66.7 mol% 1,2-dioleoyl-sn-glycero-3-phosphocholine (DOPC) and 33.3 mol% 1,2-dioleoyl-sn-glycero-3-phosphoethanolamine (DOPE) were mixed in chloroform and dried in a round-bottom flask under nitrogen flow for over 30 min. In experiments involving fluorescent lipid bilayers, 66.5% DOPC and 33.3% DOPE were mixed with 0.2 mol% 1,2-dioleoyl-sn-glycero-3-phosphoethanolamine-N-[(dipyrrometheneboron difluoride)butanoyl] (Bodipy-PE)). Dried lipid films were hydrated in HEPES buffer to a final lipid concentration of 1.0 mg/mL. The lipid solution was vortexed and underwent freeze-and-thaw cycles six times before being extruded through a 100 nm filter membrane using a mini-extruder. The LUVs were stored at 4 °C before use.

#### (b) Preparation of planar-supported lipid bilayers

Planar lipid bilayers were formed via vesicle fusion on glass. Glass coverslips were sonicated in 70% ethanol aqueous solution and then in deionized water for 15 min each, cleaned in piranha solution (3:1 H_2_SO_4_ to 30% H_2_O_2_, volume ratio) for 15 min, and then rinsed with deionized water. The etched glass coverslip was assembled into an imaging chamber. To prepare supported lipid bilayers, LUVs were diluted with HEPES buffer to a final lipid concentration of 60 μg/mL and immediately added to the imaging chamber with a pre-etched glass coverslip bottom (*31, 32*). 1.5 mM CaCl_2_ aqueous solution was added to enhance vesicle fusion on glass coverslips. After incubation at room temperature for 40 min, the formation of lipid bilayer was complete, and excessive LUVs were rinsed away with HEPES buffer thoroughly. The supported lipid bilayer was blocked with 25 μg/mL BSA for 15 min and rinsed thoroughly with dialysis buffer.

#### (c) Quantification of ISVP adsorption on supported lipid bilayers

CF568 labeled T1L/T3DM2 ISVPs were added to supported lipid bilayers at a final concentration of 10 pM (6×10^9^ ISVP/mL) at 37 °C. TIRF microscopy time-lapse images were acquired at random locations for 80 min to record the fluorescence emission of CF568. Samples were kept at 37 °C in a temperature-controlled enclosure during imaging.

### Single-virus translational and rotational imaging

#### (a) Preparation of Qdot-virus conjugates

For single-virus rotational imaging, CF568-labeled, biotinylated ISVPs were conjugated with streptavidin-coated Qdot 705. The Qdot 705 streptavidin conjugate stock solution vial was centrifuged at 5000 × g for 3 minutes before use to remove possible aggregates, following manufacturer recommended procedure. The supernatant of the Qdot 705 streptavidin conjugate was incubated with biotinylated ISVPs (2 ×10^12^ particles/mL) at a 3:1 molar ratio for 15 min. Successful conjugation was confirmed using super-resolution radial fluctuations (SRRF) imaging (*33*). The Qdot-conjugated ISVPs were used at 5 pM concentration.

ISVPs on supported lipid bilayers were imaged using TIRF microscopy for a total duration of approximately 60 s either immediately after being added to the supported lipid bilayer or after 1.5 h incubation with the supported lipid bilayer. For translational tracking, the CF568-labeled ISVPs (Em: 583 nm) were imaged at a frame rate of 6.25 Hz. For rotational tracking, CF568-labeled ISVPs and the tethered Qdot 705 (Em: 704 nm) were excited simultaneously and imaged by separating their fluorescence emissions on the same camera using a beam splitter (OptoSplit II, Cairn Research, U.K.). The beam splitter includes a 640 nm cut-off dichroic mirror to separate the 2 emission channels, a 615/40 nm emission filter for the reflected emission channel of CF568, and a 700/80 nm emission filter for the bypassed emission channel of Qdot 705. The CF568 and Qdot 705 were imaged simultaneously at an acquisition rate of 20 Hz (10). The centroids of single particles in TIRF microscopy images were localized using a radial-symmetry-based localization algorithm as previously reported (*31, 34*).

#### (b) Rotational tracking image registration

A local weighted mean mapping was used to correct the nonuniform shift between the 2 emission channels from Qdot-virus conjugates (*31, 35, 36*). Briefly, 100 nm TetraSpeck fluorescent particles (Ex/Em, 360/430 nm, 505/515 nm, 560/580 nm, 660/680 nm) were added to a coverslip at a surface density of ≈0.1 particle/μm^2^ and used as fiducial markers for image registration. By moving the motorized stage in either x- or y-direction with μm increment, the fiducial markers were imaged at each position under the same imaging condition as rotational tracking. The resulting fiducial data contained two stacks of images (at more than 100 different positions) for both channels. The fiducial markers were localized using the aforementioned radial-symmetry-based localization algorithm (*34*) and their positions in both channels were used as control points (*31*). Control points surrounding each rotational sensor were used to calculate the local weighted mean mapping, which was applied to correct the position shift of the 2 emission channels of CF568-labeled ISVP and Qdot 705 in the rotational tracking experiment. Registration inaccuracy was calculated by applying the local weighted mean mapping onto a randomly selected subset of control points. A registration inaccuracy of 6.4 nm was obtained (SI Fig. S8). The local weighted mean mapping of fiducial markers was obtained every time before the tracking of the rotational sensors.

#### (c) Detection of transient confinements and directional movements by wavelet analysis

A wavelet-based method was used to distinguish active and passive motions in the ISVP translational trajectory as previously reported (*37*). To analyze the translational dynamics, a one-dimensional Haar continuous time wavelet transform was applied to convolve the x and y displacements of an ISVP trajectory. After the transformation, the wavelet coefficients were plotted as a function of time and for different widths of the wavelet function (referred to as ‘‘scale’’). Scale 20 was chosen as a universal threshold in our wavelet analysis. If the wavelet coefficient at a certain time frame is larger than this universal threshold, the movement in these frames is active. Otherwise, the movement is passive (*38*).

#### (d) Calculation of mean-square displacements (MSD)

The MSD was calculated as a function of lag time *τ* based on the following equation:

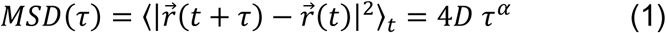

where *r⃗* is the position of the ISVP, *r⃗*(*t* + *τ*) − *r⃗*(*t*) is the end-to-end displacement of a single ISVP from time point *t* to time point (*t* + *τ*), 〈 〉 means average over time, *D* is the diffusion coefficient, and *α* is the anomalous diffusion exponent averaged over the entire trajectory of a single ISVP (*31, 39, 40*). *α* was obtained from fitting the log−log MSD plots using the following equation:

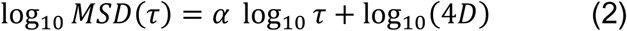

*α* carries information about the local motion type: for *α* =1, the ISVP undergoes Brownian diffusion; for *α* <1, the local motion is subdiffusive; whereas for *α* >1, the local motion is superdiffusive (*41*).

#### (e) Calculation of turning angle

The turning angle *β* during ISVP diffusion was calculated based on the displacement angle relative to the preceding step over the specified time interval (*40*):

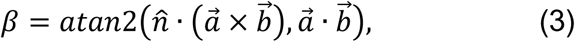

where *a⃗* and *b⃗* are adjacent steps, and *n̂* is the unit normal vector to the membrane plane, which sets the sign convention for the turning angle. When viewed along +*n̂*, positive angles indicate a direction counterclockwise to the previous step, and negative angles indicate a direction clockwise to the previous step.

#### (f) Calculation of the probability distribution of displacement

The probability distribution of ISVP displacement, *P*(|*R*(*t*) − *R*(*t* = 0)| > *r*), represents the probability of ISVP displacement at time t (*R*(*t*)) relative to that at time 0 (*R*(*t* = 0)) is larger than a given distance *r*. The probability was then plotted against *r* (*42*).

#### (g) Calculation of the persistence length of a trajectory

The translational trajectory of ISVP can be regarded as a polymer chain, and the persistence length can be calculated to evaluate the linearity of the trajectory(*43*):

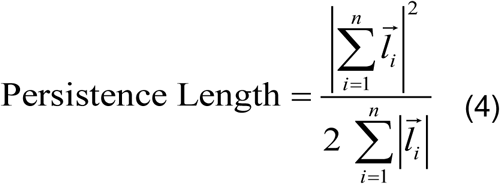

Where 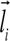 is a single step, 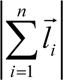 is the net displacement (end-to-end distance), and 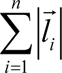 is the total displacement (contour length).

#### (h) Rotational tracking analysis

After rotational tracking and image registration, the centroids of the projection of ISVP (*x*_*ISVP*_, *y*_*ISVP*_) and Qdot (*x*_*Qdot*_, *y*_*Qdot*_)on the same rotational sensor in each frame were connected as a vector (*x*_*Qdot*_ − *x*_*ISVP*_, *y*_*Qdot*_ − *y*_*ISVP*_), of which the in-plane angle (orientation) was obtained as *φ*. The length of the vector was used to calculate the out-of-plane angle, *θ*, using the equation below:

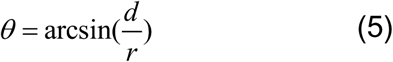

where *d* is the projected interparticle distance and *r* is the physical interparticle distance (*31*). Because *r* varies slightly among ISVPs due to particle size distribution, it was measured from microscopy images as the maximal projected interparticle distance *d*_*max*_, which corresponds to the largest diameter observed as the ISVP rotates through different orientations on the membrane. The lower bound of *r* was set to 50 nm, considering the average radii of ISVP and streptavidin-coated Qdot 705 (*31, 44*).

### Computational model

#### (a) Overview of the model

We developed a stochastic computational model to investigate possible mechanisms underlying the observed motion of individual ISVPs on a planar supported lipid bilayer. In the model, a particle is in contact with a two-dimensional surface representing the lipid bilayer. The dynamics are described by a set of possible actions and associated rates that are informed by the underlying biology and experimental observations. We used the Gillespie algorithm (*45, 46*) to generate stochastic simulation trajectories and investigate particle motion across an ensemble of independent particles.

In the model, the particle has discrete sites containing cleavable “peptides” distributed around its surface. These sites represent points of contact between the particle and the surface, and when a peptide is present, it can be “inserted” into the membrane. When a peptide is inserted, the viral particle diffuses with a bias in a randomly chosen direction. This preferred direction is maintained until a different neighboring peptide is inserted, the particle rotates to a site without a peptide, or the peptide is cleaved. When a site without a peptide is in contact with the surface, the viral particle undergoes isotropic diffusion.

When cleaved, a peptide remains inserted in the bilayer, where it is represented as a particle that diffuses in two dimensions. Inserted peptides can aggregate to form larger clusters, which are also represented by diffusing particles. The peptide clusters attract the viral particle at short ranges, with larger clusters exhibiting stronger attraction and slower diffusion.

#### (b) ISVP model

The ISVP is modeled as a sphere with 200 sites distributed on the surface with neighboring sites approximately equidistant from one another. Each site initially contains a “peptide,” which physically corresponds to a small number of μ1N peptides that can be inserted into the membrane. In the model, only one site is in contact with the surface at a time. If the site contains a peptide, it is assumed to be inserted in the membrane. The particle can rotate to neighboring sites: The rate is 0.002 *τ*_0_^−1^ per site containing a peptide and 0.00025 *τ*_0_^−1^per empty site. Here, *τ*_0_denotes the simulation time unit. When a peptide is inserted, it is cleaved at a rate of 0.01 *τ*_0_^−1^. When a peptide is cleaved, it initially resides in the membrane at the location of the viral particle, and its site on the viral particle remains empty for the remainder of the simulation. The simulation begins with the ISVP in contact with the membrane surface with a peptide inserted.

#### (c) Peptide model

After it is cleaved, a peptide undergoes diffusion in the membrane and can aggregate with other cleaved peptides (*47*). Each peptide cluster is represented as a particle that is characterized by the number of peptides in the cluster; a single peptide is considered a cluster of size 1. Aggregation occurs when two peptide clusters are in close proximity. When they are < 20 *σ* apart, a new particle replaces the previous two (*σ* denotes the simulation distance unit). The new cluster contains all of the peptides in the previous two, with its initial position at the average of the original two weighted by the number in each cluster. The maximum cluster size is 11 peptides, so if two clusters have more than 11 peptides in total, they cannot aggregate.

We model the attraction of the ISVP to peptide clusters using a truncated and shifted 9-6 Lennard-Jones potential(*48, 49*) with a maximum interaction distance of 20 *σ*. This imposes a short-range repulsion, preventing overlap, with a longer-range attraction. The strength of the interaction increases linearly with the number of peptides in the cluster.

#### (d) Lateral motion

Lateral motion occurs via a stochastic hopping process. An isolated viral particle undergoes isotropic diffusion when no peptide is inserted, hopping a distance of 1 *σ* in a randomly chosen direction at a rate of 1 *τ*_0_^−1^. When anchored to the surface by a peptide, it undergoes biased motion imposed by a direction-dependent acceptance-rejection criterion. A trial hop is generated as for isotropic diffusion and is accepted with probability exp(−2*θ*), where *θ* is the smallest angle between the preferred direction and the attempted hop direction. Otherwise, the trial move is rejected.

When peptides are embedded in the membrane, the attraction between the viral particle and peptide clusters imposes additional constraints on the viral motion. For each of the lateral moves described above (isotropic and biased), the move is accepted if the change in energy is ≤ 0. In contrast, the move is accepted with probability exp(−Δ*E*) when the move is energetically unfavorable (Δ*E* > 0).

Peptide clusters undergo isotropic diffusion. A peptide cluster of size *N*_*c*_ hops a distance of 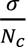 in a randomly chosen direction at rate 1 *τ*_0_^−1^. With this choice, the diffusion coefficient of a peptide cluster decreases quadratically with the number of peptides.

### Statistical analysis

All data are presented as mean ± S.E.M. Statistical evaluations of data were performed by Student’s t-test for two groups and one-way analysis of variance for multiple groups. Statistical significance is indicated as follows: ****p < 0.0001, ***p < 0.001, **p < 0.01, *p <0.05, and not significant (NS) p > 0.05. Statistical figures were plotted using OriginLab Pro 2023b.

## RESULTS

### μ1 peptide release accelerates ISVP adsorption on lipid membranes

We first investigated ISVP adsorption on a planar lipid membrane using single-virus imaging. Supported bilayers composed of 66.7 mol% 1,2-dioleoyl-sn-glycero-3-phosphocholine (DOPC) and 33.3 mol% 1,2-dioleoyl-sn-glycero-3-phosphoethanolamine (DOPE) were used. The 2:1 molar ratio mirrors the lipid composition of endosomal membranes and facilitates ISVP-to-ISVP* conversion (*7, 8*) (Fig. 1C). Fluorescently labeled Type 1 Lang (T1L) ISVPs (10 pM) were added and their binding to the lipid bilayer was monitored over 80 min at 37°C using time-lapse Total Internal Reflection Fluorescence (TIRF) microscopy (Fig. 2A).

**Figure 2.**
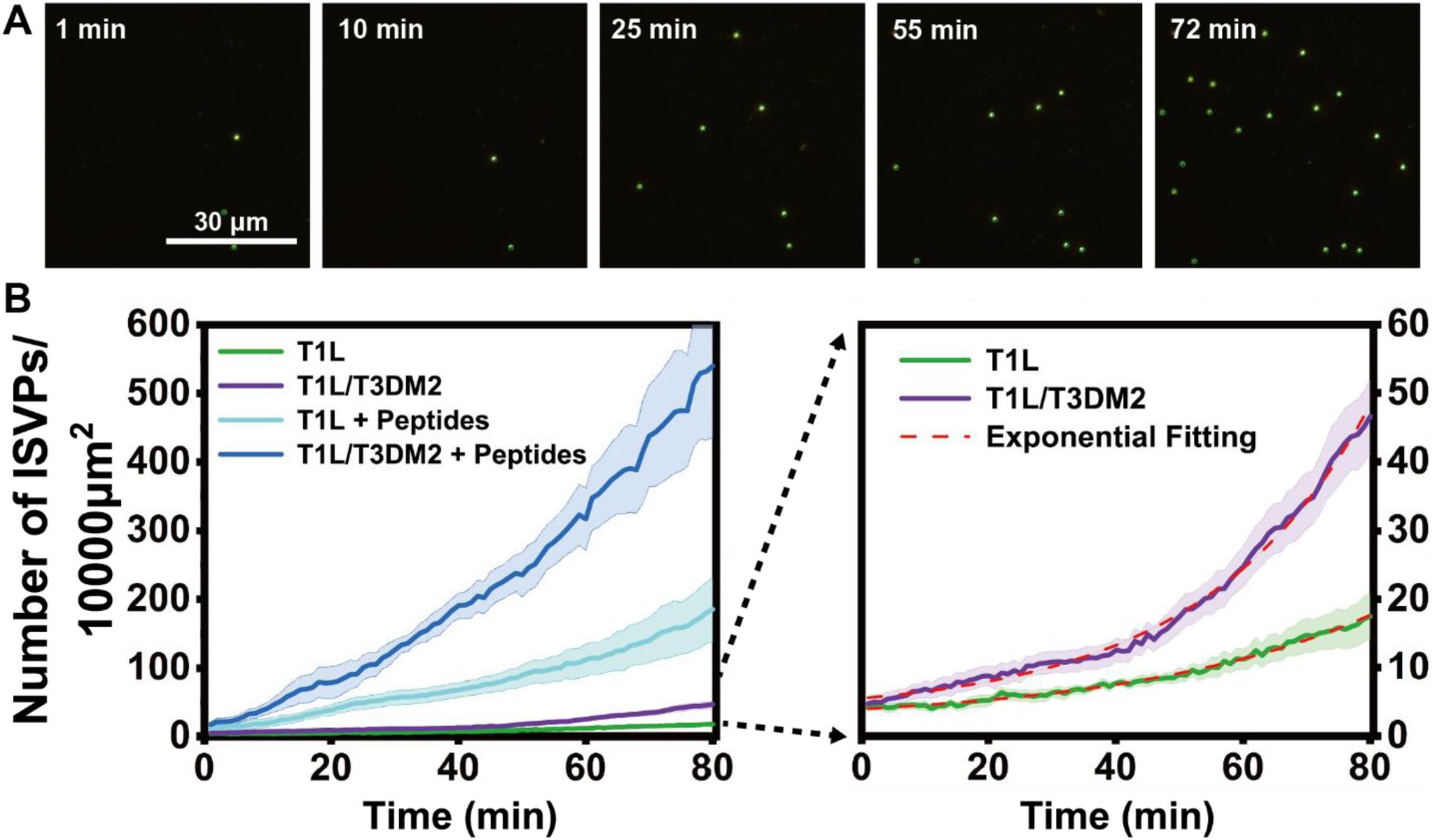
Recruitment of reovirus ISVPs on the planar-supported lipid bilayer. (A) Time-lapse TIRF Fluorescence microscopy images showing the recruitment of T1L/T3DM2 ISVPs on the planar-supported lipid bilayer at the same location over time. (B) Averaged line plots showing the adsorption of 10 pM of T1L and T1L/T3DM2 ISVPs with or without the presence of excessive peptides produced *in vitro* on the supported lipid bilayer over time. The right figure is the y-axis zoom-in view of the left. Error bars represent the standard error of the mean (S.E.M.).

We observed that the surface density of adsorbed ISVPs increased exponentially over time (Fig. 2B), indicating that ISVP adsorption promotes the binding of additional ISVPs. One possible explanation for this observation is that adsorbed ISVPs directly attract others. However, this is unlikely, because ISVP aggregation requires much higher concentrations and no aggregation was observed under our experimental conditions (Fig. 2A). A more plausible explanation is based on our previous findings that ISVP interaction with lipid membranes under physiological conditions promotes conversion of ISVPs to ISVP*s, which triggers release of peptides from the N- and C-terminus of the μ1 protein (*4, 5, 7*). Among these, the N-terminal peptide μ1N enhances adsorption of additional ISVPs (Fig. 1B).

To test this idea, we compared adsorption of T1L ISVPs to that of reassortant strain T1L/T3DM2. T1L and T1L/T3DM2 reovirus differ only in the origin of their μ1-encoding M2 gene segment (*10, 25*) and both encode an identical μ1N peptide sequence. The key distinction is that T1L/T3DM2 undergoes ISVP conversion more rapidly, leading to faster and more efficient release of μ1-derived peptides, including μ1N (*25*). Owing to this difference, T1L/T3DM2 initiates infection at a faster rate than T1L (*50*). We found that, like T1L ISVPs, T1L/T3DM2 ISVPs also followed exponential adsorption kinetics, but adsorbed at a faster rate. These data are consistent with the more efficient uncoating and release of μ1N by T1L/T3DM2 (Fig. 2B). However, it is possible that the regions of μ1 that differ between the T1L and T3D μ1 proteins (15 of 708 amino acids) affect the surface properties of ISVPs and regulate the μ1N-mediated adsorption of ISVPs to membranes. To test this possibility, we used the T1L/T3DM2 D371A mutant, a strain that maintains ISVP surface properties comparable to T1L/T3DM2 but is impaired in μ1N release (*51*). Adsorption of D371A mutant followed a linear, rather than exponential, adsorption trend (Fig. S1). These results indicate that the μ1N peptide release is crucial for the accelerated increase in ISVP adsorption. To further validate that increased ISVP adsorption was mediated by μ1N, we pretreated the supported lipid bilayer with ISVP* supernatant, a solution containing μ1N, before ISVP introduction. We found that this pre-treatment further increased the adsorption rate of both T1L and T1L/T3DM2 by more than 10-fold (Fig. 2B). Together, our data support the model for positive feedback where ISVP interaction with lipid membranes promotes the release of μ1N, which in turn recruits additional ISVPs to the membrane.

### μ1N peptide release affects persistence and confinement of ISVP dynamics

While the above data suggest the released μ1N promotes ISVP interaction with the lipid membrane, they do not reveal details of the events occurring at the particle-membrane interface. To obtain a better understanding of this process, we investigated the motion characteristics of T1L and T1L/T3DM2 ISVPs on lipid membranes by tracking the individual trajectories of ISVPs for 60s immediately after their addition and after 1.5 h of interaction (Fig. 1A). The 60s observation period was chosen to capture the early particle-membrane interactions while minimizing photobleaching effects (Fig. S2). The trajectories showed that T1L ISVPs exhibited more directional segments of motion initially, which transitioned to confined ones after prolonged membrane interaction. This also resulted in a decreased range of motion with time. In contrast, T1L/T3DM2 ISVPs started with relatively confined segments of motion and remained predominantly confined throughout the interaction (Fig. 3A).

**Figure 3.**
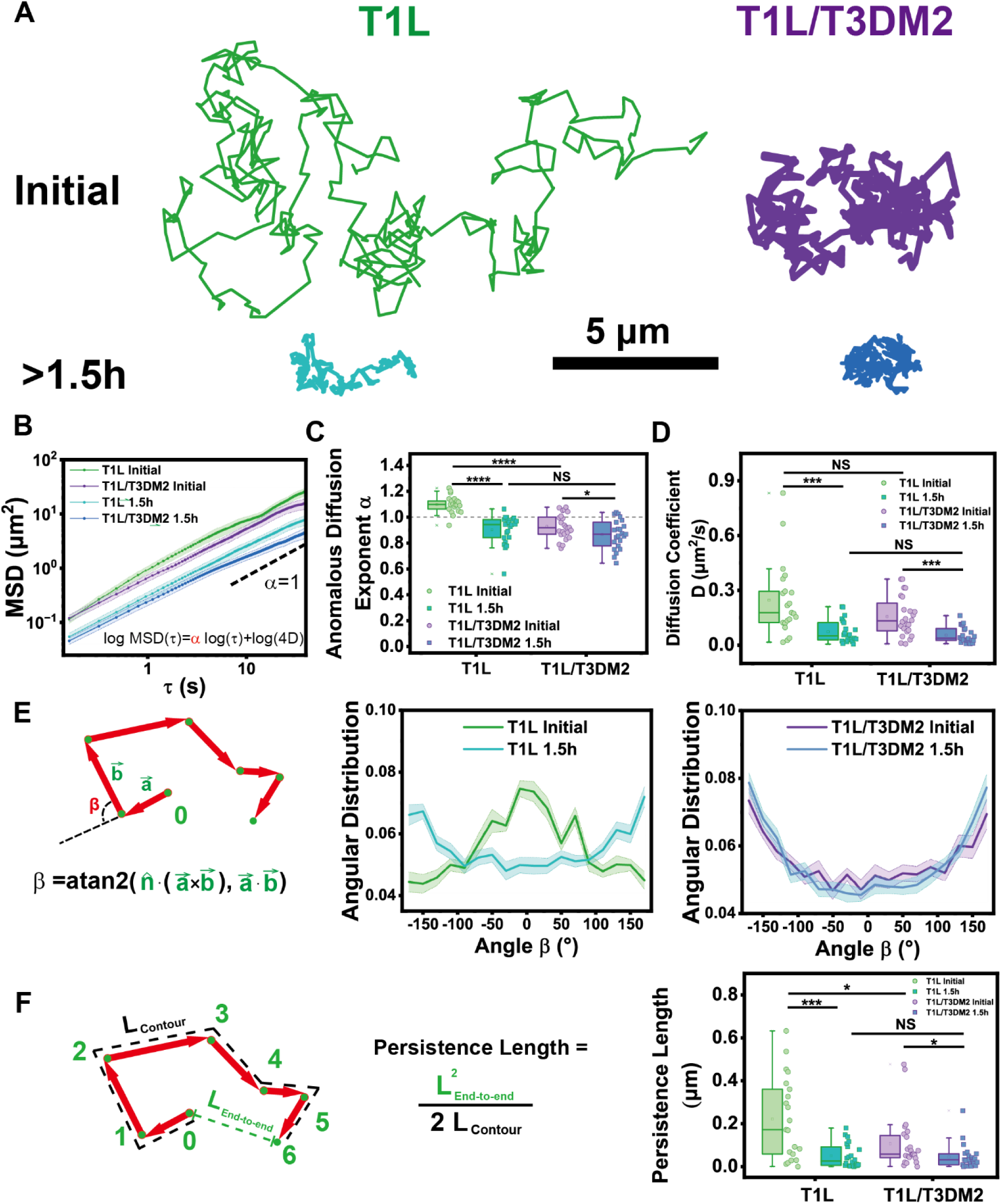
Translational diffusion of ISVPs on the planar-supported lipid bilayer. (A) Representative line plots showing the trajectories of T1L and T1L/T3DM2ISVPs on the supported lipid bilayer. Scale bar, 5 μm. (B) Averaged log-log line plots showing the mean square displacement of ISVP trajectories. T1L initial (n=21), T1L/T3DM2 initial (n=24), T1L after 1.5h (n=22), and T1L/T3DM2 after 1.5h (n=23). Error bar represents the standard error of the mean (S.E.M.). (C and D) Statistical analysis of anomalous diffusion exponent (C) and diffusion coefficient (D) of trajectories from T1L initial (n=21), T1L/T3DM2 initial (n=24), T1L after 1.5h (n=22), and T1L/T3DM2 after 1.5h (n=23). (E) Schematic illustration of the turning angle β, the change of angle between two consecutive steps in a single trajectory, and averaged line plots showing the distribution of turning angle β for trajectories of T1L (n=22) and T1L/T3DM2 (n=21) upon the initial adsorption on bilayers. Error bar represents the standard error of the mean. (F) Schematic illustration of the persistence length in a trajectory and statistical analysis of persistence length of trajectories from T1L initial (n=21), T1L/T3DM2 initial (n=24), T1L after 1.5h (n=22), and T1L/T3DM2 after 1.5h (n=23). Each boxplot indicates the interquartile range from 25% to 75% of the corresponding data set. The mean and median are demonstrated as the square and the horizontal line, respectively. Statistical significance is highlighted by p-values.

This dynamic characteristic was quantitatively confirmed using mean squared displacement (MSD) analysis (Fig. 3B). MSD for two-dimensional diffusion follows the power-law relationship: *MSD* (*τ*) ∝ *Dτ*^*α*^, where *τ* is time, *D* is the effective diffusion coefficient, and *α* is the anomalous diffusion exponent. This exponent *α*, extracted as the slope from log-log plots of MSD(*τ*) versus *τ*, indicates the type of diffusion: Brownian (*α* = 1), directional (*α* > 1), or confined (*α* < 1). T1L ISVPs exhibited directional motion (*α* > 1) initially and transitioned to confined motion (*α* < 1) after prolonged interaction (Fig. 3C). This transition correlated with a significant decrease in diffusion coefficient (Fig. 3D). In contrast, T1L/T3DM2 ISVPs displayed confined motion (*α* < 1) from the outset, with a portion immobilized on the bilayer during the initial imaging (Fig. S3). Over time, confinement increased further (Fig. S4A, B), as indicated by decreases in both diffusion exponent *α* and diffusion coefficient *D* after 1.5h of lipid interaction for both strains (Fig. 3C, 3D). This progressive confinement was also confirmed by plotting the probability of displacements over time, where reduced displacement probabilities indicated smaller trajectory sizes (Fig. S5A-G).

To quantitatively confirm the directionality of ISVP motion, we analyzed the turning angle (*β*) between consecutive steps in each trajectory (*40*) (Fig. 3E). The range of turning angle spans from −180° to 180°. For T1L ISVPs, turning-angle distributions were initially centered near 0°, indicating directional persistence. After 1.5h, the turning angle distribution inverted and shifted toward ±180°, indicating retrograde movement, which is consistent with the transition to more confined behavior. In contrast, the turning angle of T1L/T3DM2 ISVPs peaked consistently at (±180°) throughout both phases, suggesting more confined movement overall, with less inclination toward directional motion.

To quantify the long-range directional persistence of ISVP motion, we computed the persistence length by modeling each trajectory as a flexible polymer chain (*43*). In a linear polymer, the persistence length marks the contour length below which the chain behaves as a rigid rod, i.e., its local tangents are correlated. Analogously, the persistence length of a particle trajectory reflects the scale over which directional motion is maintained before reorientation. During the initial phase of interaction, T1L ISVPs exhibited an average persistence length of ≈ 222 nm, roughly three times the diameter of an ISVP particle (70–80 nm) (Fig. 3F). After 1.5 h of interaction, this length decreased significantly to ≈ 51 nm, indicating reduced directional movement. In contrast, T1L/T3DM2 ISVPs displayed shorter persistence lengths throughout the interaction, with a slight decrease over time, but both initial and later values were notably smaller than those of T1L ISVPs.

The results together indicate that T1L ISVPs initially exhibited predominantly directional movement after adsorption, which gradually transitioned to confined motion. In contrast, T1L/T3DM2 ISVPs displayed confined movement from the start, becoming progressively more restricted over time. These findings suggest that the rate of μ1N peptide release significantly influences the non-Brownian dynamics of both T1L and T1L/T3DM2 ISVPs.

### ISVPs move directionally between confinements at discrete binding sites

We next sought to determine the basis of what drives the directional motion of T1L ISVPs. To this end, we examined the single trajectories of ISVPs to assess how they moved using wavelet transform analysis. Wavelet analysis segments each trajectory into short time windows and detects brief changes in trajectories by comparing local motion to simple templates, allowing us to quantify transient and heterogeneous dynamical changes (*37, 38*). The ensemble-averaged analyses above revealed that T1L ISVPs exhibited directional movement during the early stage. However, with wavelet analysis, we discovered that the motion was not entirely directional; rather, it was a mixture of directional jumps and some confined behavior (Fig. 4A). Surprisingly, even in the confined motion of both T1L and T1L/T3DM2 ISVPs after prolonged membrane interaction, there were small percentages of directional jumps, with the majority of the motion still confined (Fig. 4C).

**Figure 4.**
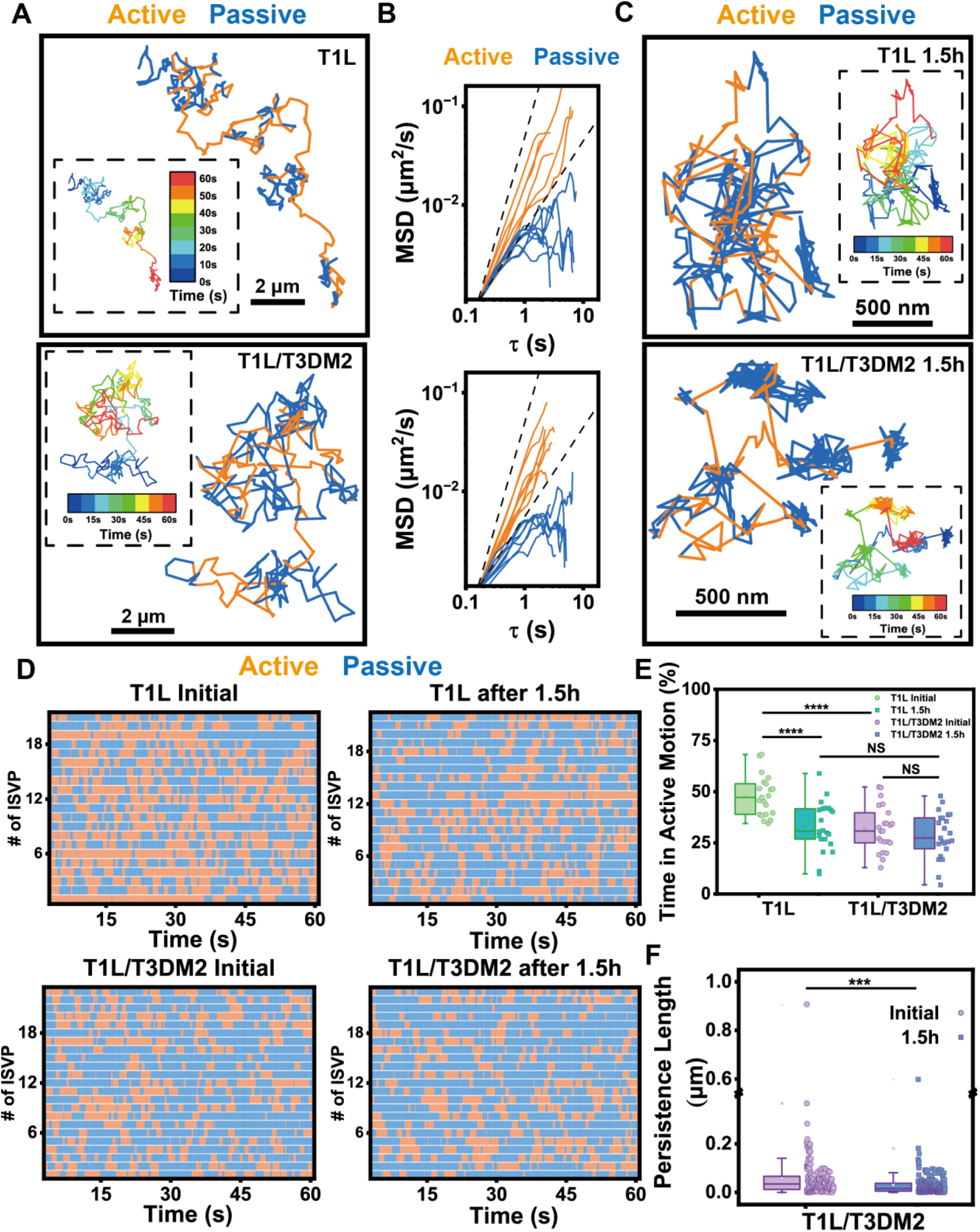
Active and passive segments within ISVP translational trajectories. (A) Representative line plots showing the trajectories of T1L and T1L/T3DM2 ISVPs on the planar-supported lipid bilayer right after the recruitment to the membrane. Scale bar, 2 μm. (B) Log-log line plots showing the mean square displacement of active and passive segments from T1L ISVPs and T1L/T3DM2 ISVPs right after the recruitment to the planar-supported lipid bilayer. (C) Representative line plots showing the trajectories of T1L and T1L/T3DM2 ISVPs on supported lipid bilayer 1.5h after incubation with the planar-supported lipid bilayer. Scale bar, 2 μm. (D) Color map showing the active and passive segments within trajectories of ISVPs on supported lipid bilayer from T1L initial (n=21), T1L/T3DM2 initial (n=24), T1L after 1.5h (n=22), and T1L/T3DM2 after 1.5h (n=23). (E) Statistical analysis of the percentage of time in active motion within trajectories from T1L initial (n=21), T1L/T3DM2 initial (n=24), T1L after 1.5h (n=22), and T1L/T3DM2 after 1.5h (n=23). (F) Statistical analysis of persistence length of passive diffusional segments from T1L/T3DM2 initial (n=172), and T1L/T3DM2 after 1.5h (n=185). Each boxplot indicates the interquartile range from 25% to 75% of the corresponding data set. The mean and median are demonstrated as the square and the horizontal line, respectively. Statistical significance is highlighted by p values.

A crucial part of the wavelet analysis is to distinguish active and passive segments of individual ISVP trajectories. Active segments, corresponding to directional jumps, were characterized by an anomalous diffusion exponent (*α* > 1), whereas passive segments, indicative of either Brownian or confined motion, had *α* values close to or less than 1 (Fig. 4B). Wavelet-based segmentation resolved alternating directional long jumps, which allowed the particles to transition to a different location and confined states within single trajectories (Fig. 4A-C). Both strains exhibited this two-state behavior, while the number and length of these “jumps” varied between the two ISVPs and across different interaction phases (Fig. 4D, E). Specifically, during the initial interaction phase, T1L ISVPs spent half of their time undergoing directional movements. The duration of these directional movements decreased over time, reflecting increased local confinement. For T1L/T3DM2 ISVPs, they exhibited predominantly confined motion with some directional jumps, but the scale of confinement further increased after prolonged membrane interaction, as indicated by the decreased persistence length of the passive segments, suggesting progressively stronger ISVP-lipid interactions over time (Fig. 4F).

Interestingly, during the later phase of interaction, both ISVPs occasionally returned to previously visited confinement sites (Fig. 4C). This behavior suggests that ISVPs are attracted to specific, discrete sites on the lipid membrane, with this attraction intensifying over time. These sites are likely formed by the collection of μ1N peptides, which are released and inserted into the membrane, and create localized points of attraction for the ISVPs (Fig. 1C). But what drives ISVPs to jump between these binding sites instead of remaining at a single site?

We hypothesized that the dissociation of μ1N peptides from the ISVP capsid drives the dynamic jumps of ISVPs. This was supported by observations from ISVP rotational tracking experiments. Using streptavidin-labeled Qdot 705 attached to biotinylated ISVPs, we imaged and tracked the rotation of single ISVPs during their translational movement on lipid membranes (Fig. 5A). Right after adsorption onto planar-supported lipid bilayers, ISVPs present a combination of active directional movements and passive confined movements (Fig. 5B). In contrast, the T1L/T3DM2 D371A mutant, which does not release μ1N peptides, remained stationary on the lipid bilayer (Fig. S6 A-G), showing only random orientational fluctuations due to thermal motion (Fig. S7A, B). This random thermal motion is different from the μ1N peptide aggregation-induced transient confinement zones (Fig. 4C, 5C). A representative T1L/T3DM2 ISVP, with higher kinetics of μ1N release, exhibited ordered rotational movement with slight in-plane orientation changes (Fig. 5C, S7B). Moreover, the ISVP transitioned back and forth between three adjacent binding sites (Fig. 5C). Our rotational analysis not only confirmed directional movements but also distinguished the differences between transient confinement zones and the thermal motion despite the similar translational subdiffusion pattern.

**Figure 5.**
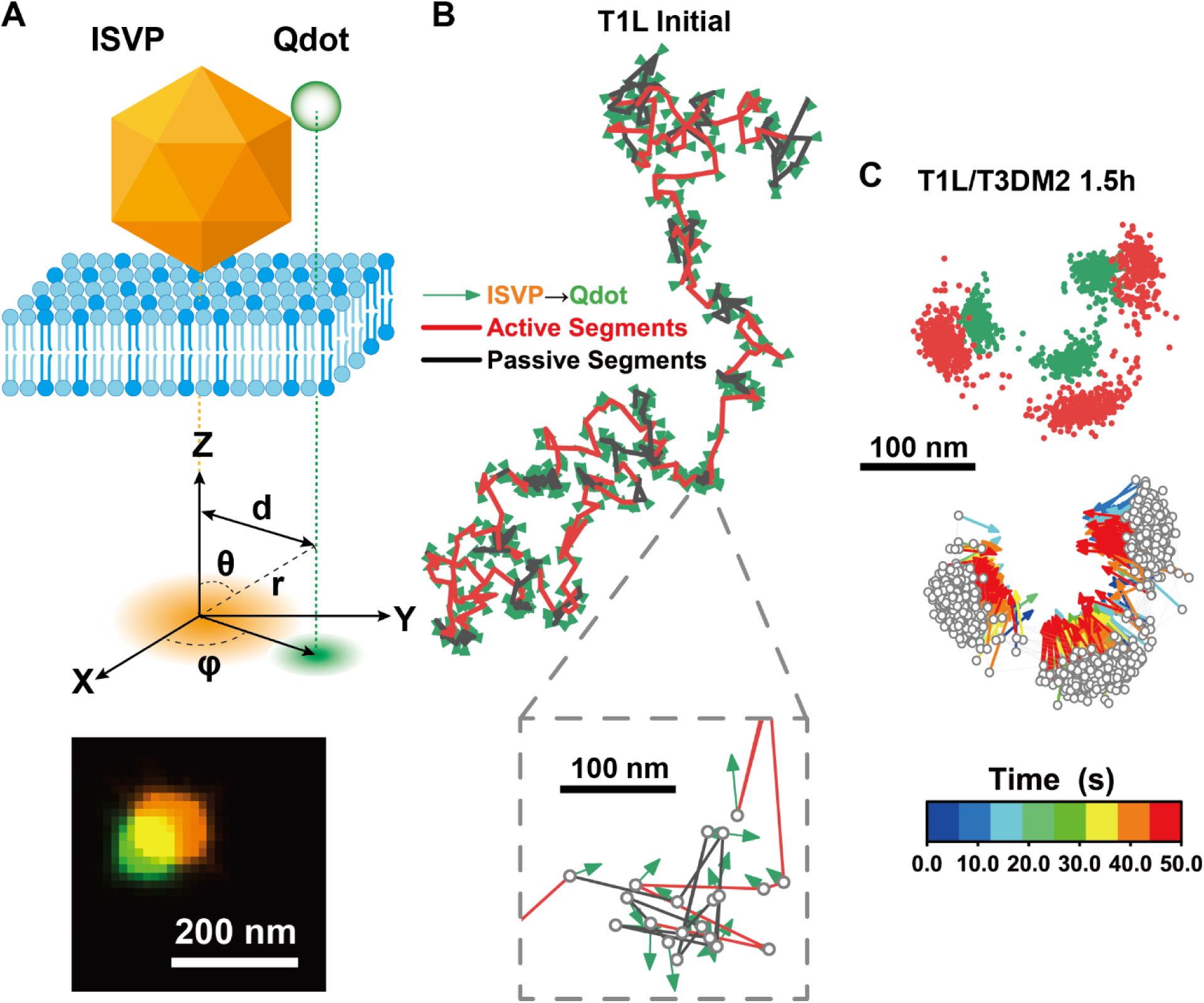
Rotational tracking of ISVPs on the planar-supported lipid bilayer. (A) Schematic illustration of rotational tracking of ISVP diffusion on the planar-supported lipid bilayer. (B, C) Representative rotational trajectories of T1L (B) and T1L/T3DM2 (C) ISVPs on the planar-supported lipid bilayer. Colormap encodes temporal information.

### Computer simulations suggest peptide clusters promote confined motion

Based on our experimental observations, we next developed a stochastic dynamical model to capture essential features of the ISVP motion on a 2D surface and understand the complex interactions governing such dynamics (Fig. 6A). In the model, the ISVP undergoes biased motion when a capsid-anchored peptide is inserted in the membrane and unbiased diffusion otherwise. When peptides are released from the ISVP surface, they diffuse in the plane of the membrane, aggregate to form slow-moving peptide clusters, and attract the ISVP at short range. The model allowed us to mimic both T1L and T1L/T3DM2 ISVPs by varying the release rate of peptides from the capsid, with T1L corresponding to slow release of peptides and T1L/T3DM2 corresponding to fast release. For each case, we analyzed ISVP motion over many independent simulation trajectories. In contrast with experiments, simulations allowed both the ISVP and cleaved peptides to be tracked simultaneously.

**Figure 6.**
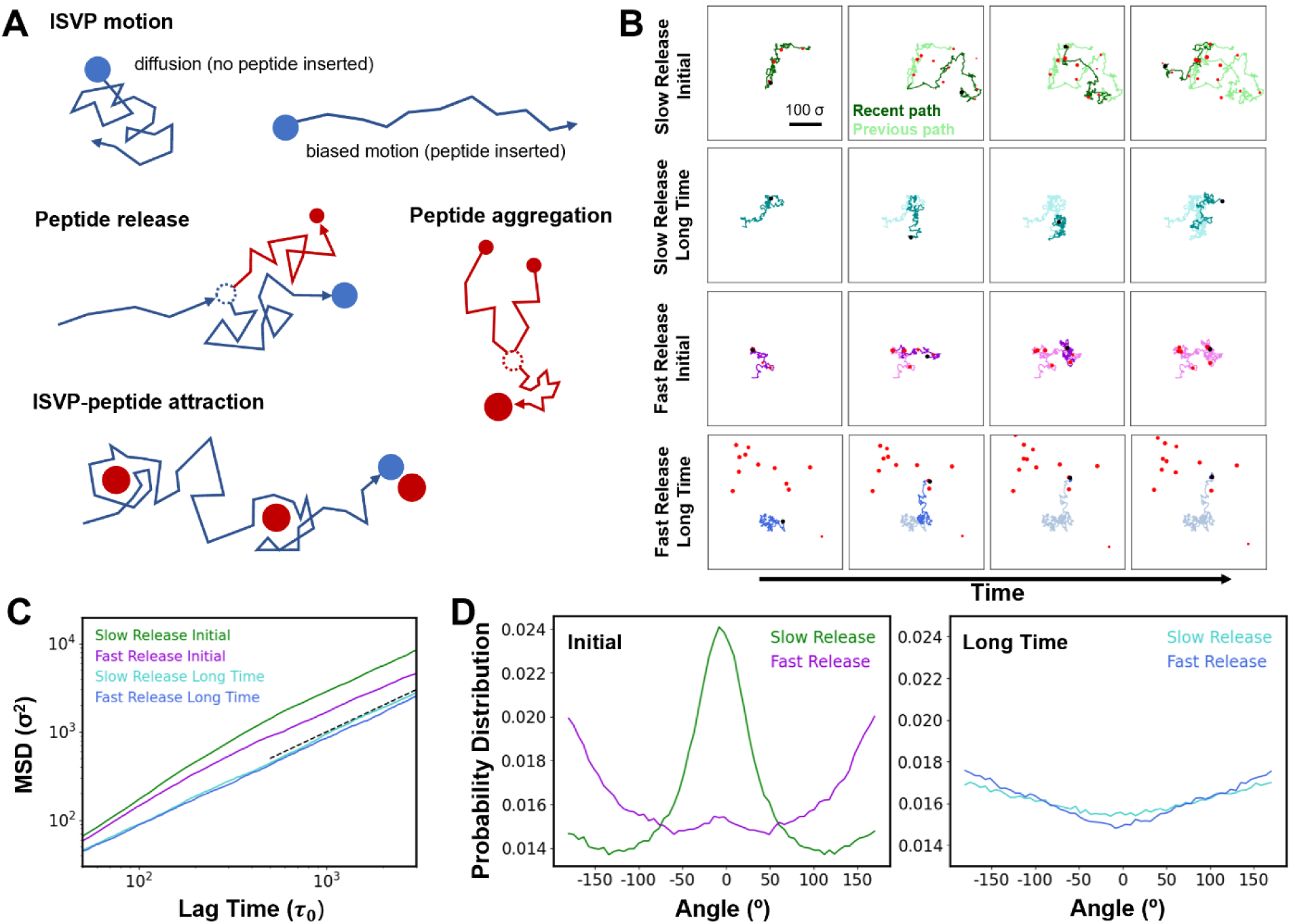
Computational model of ISVP motion. (A) Schematic representation of the model. The ISVP (blue circle) undergoes stochastic motion on a 2D membrane surface. Characteristics of the motion depend on the state of the system. Peptides (red circles) are cleaved from the ISVP surface, diffuse in the plane of the membrane, aggregate to form peptide clusters (larger red circles), and attract the ISVP at short range. (B) Snapshots from four representative simulation trajectories. Each row is a different case sampled at increasing time from left to right. Trajectories are shown at intervals of 6,000 *τ*_0_. The most recent part of the trajectory is shown in a darker shade compared to previous segments. Long-time results start at 9 × 10^5^ *τ*_0_, when most peptides are cleaved on average. Peptide clusters are depicted by red clusters. (C) Mean square displacement (MSD) from simulations. The dashed black line has a slope of 1. (D) Distribution of turning angle from simulations. Results are shown for initial time (left) and long time (right). Angles were determined at a sampling time of 100 *τ*_0_. The MSD and distributions were determined from 5,400 independent trajectories.

Time traces of individual trajectories revealed differences between ISVPs with slow and fast peptide release (Fig. 6B). With slow release, as in the case of T1L ISVP, particle motion was characterized by localized diffusion interspersed with longer, relatively straight paths. Because of the slow peptide release rate, peptides were deposited far apart in the membrane, suppressing their aggregation into large clusters. As time progressed, the ISVP became transiently associated with some peptide clusters, but it was able to explore a relatively large area. At long times, because the peptides did not form large aggregates, they dispersed throughout the membrane and did not hinder the ISVP motion. In this regime, the ISVP motion was diffusive. In contrast, for the case of T1L/T3DM2 with fast peptide release, the peptides were inserted into the membrane much closer to one another, promoting aggregation into large, relatively immobile clusters. The large clusters served as localized traps that attracted the ISVP at both short and long times, suppressing its motion.

The analysis of many independent simulation trajectories revealed that when the ISVP was initially in contact with the surface, the MSD exhibited directional (*α* > 1) motion at short times, with the directional motion most pronounced for slow peptide release (Fig. 6C). At long times, ISVPs with both slow and fast peptide release exhibited slower motion with *α* close to 1. Differences between slow and fast peptide release were also reflected in the distribution of angles between neighboring segments of ISVP traces (Fig. 6D). Initially, slow peptide cleavage produced a marked peak centered at β = 0°, indicating directed motion. In contrast, the peak was strongly suppressed for fast cleavage, with much more weight near ±180°, indicating the influence of confinement by peptide clusters. At long time, both of the distributions were relatively flat, with the increased weight at large angles arising from confined motion near peptide clusters.

In the model, insertion of a capsid-associated peptide into the membrane generates directed motion on timescales associated with peptide release. Once released and inserted into the membrane, peptides can locally trap an ISVP due to their mutual attraction. Faster peptide cleavage promotes the formation of larger peptide clusters because peptides are cleaved in closer proximity and can more readily aggregate. Further, if the ISVP becomes locally trapped by a peptide cluster, subsequent release events provide more peptides that can join the cluster, resulting in positive feedback. The simulation results capture the dynamics of T1L and T1L/T3DM2 ISVPs, and suggest a process by which the processive three-way interactions between the ISVPs, peptides, and lipid membranes drive the heterogeneous motion of individual ISVPs.

## Discussion

This study sheds light on how non-enveloped reovirus particles interact with lipid membranes by directly imaging and quantifying single-virus motion. Using a planar-supported lipid bilayer model system and reovirus strains with distinct uncoating kinetics, we addressed critical questions regarding virus-lipid interactions and the mechanisms driving adsorption, diffusion, and endosomal escape.

Our findings demonstrate that the μ1N penetration peptide plays a pivotal role in enhancing virus binding to the membrane. The membrane-inserted μ1N, released during the ISVP-to-ISVP* conversion, creates transient binding sites that promote adsorption of additional viruses. This self-propagating mechanism highlights the role of μ1N peptides in amplifying the recruitment of viral particles to the lipid bilayer.

Through single-particle tracking experiments, we uncovered how the release of μ1N peptides propels ISVP diffusion on membranes. During uncoating, the dissociation of μ1N peptides from the capsid breaks the bond between the ISVP capsid and lipid membrane. Our computational model, which captures the complex dynamics observed in experiments, suggests directional movements in ISVP trajectories associated with μ1N peptide insertion in the membrane. This dynamic behavior exemplifies how capsid uncoating can drive viral diffusion.

We have previously shown that the ISVP-to-ISVP* conversion is facilitated by the lipid bilayer (*7*). The amphiphilic μ1N peptides released during this process act as binding sites within the membrane, enhancing ISVP-membrane interactions. Over time, these peptides may aggregate, forming molecular anchors where ISVPs adsorb and become temporarily trapped before escaping via thermal fluctuations (*47*). In addition to the membrane permeability induced by the peptide insertion, the directional diffusion driven by ISVP capsid uncoating allows the ISVP to move around an area larger than a peptide-induced pore, release additional peptides and further destabilize the membrane. This mechanism could progressively enlarge these pores to facilitate viral penetration and endosomal escape. As we previously reported in a GUV model system (*10*), even a single virus can exert substantial effects on membrane integrity, underscoring the profound impact of this mechanism on endosomal escape, particularly in the confined space of an endosome.

In summary, this study provides quantitative evidence revealing how reoviruses orchestrate a self-propagating mechanism to achieve membrane penetration and initiate infection. Our findings highlight a process where virus-membrane interactions begin without specific high-affinity ligand-receptor binding but evolve into a system where the virus itself generates membrane binding sites, amplifying its interaction with the membrane. As reovirus serves as a model non-enveloped virus, these insights likely extend to other non-enveloped viruses, offering a broader understanding of their infection mechanisms and informing strategies to combat viral diseases.

## AUTHOR CONTRIBUTIONS

M.J. and Y.Y. designed research; M.J., G.R.C., Y.-q.Y., A.J.S., S.M.A., and P.D. performed research; M.J., G.R.C., and S.M.A. analyzed data; and M.J., S.M.A., P.D., and Y.Y. wrote the paper.

## DATA AVAILABILITY

All data are included in the manuscript and/or supporting information. Materials will be shared upon request.

## Supplementary Information for

### Contents

Supplementary figures S1-S8

**Figure S1.**
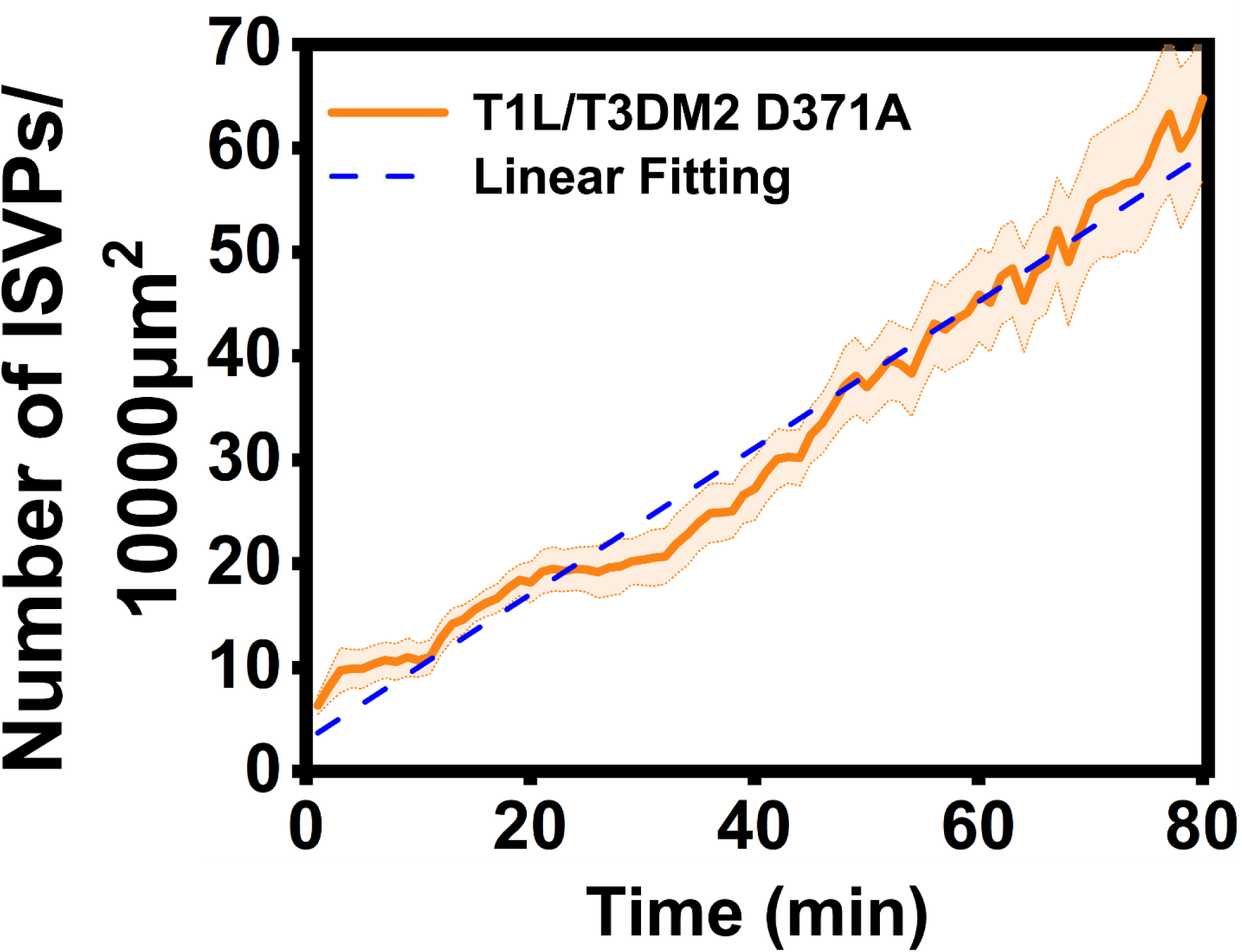
Recruitment of mutant reovirus ISVPs on the planar-supported lipid bilayer. Averaged line plots showing the adsorption of 10 pM T1L/T3DM2 D371A ISVPs on supported lipid bilayer along time. Error bar represents the standard error of the mean (S.E.M).

**Figure S2.**
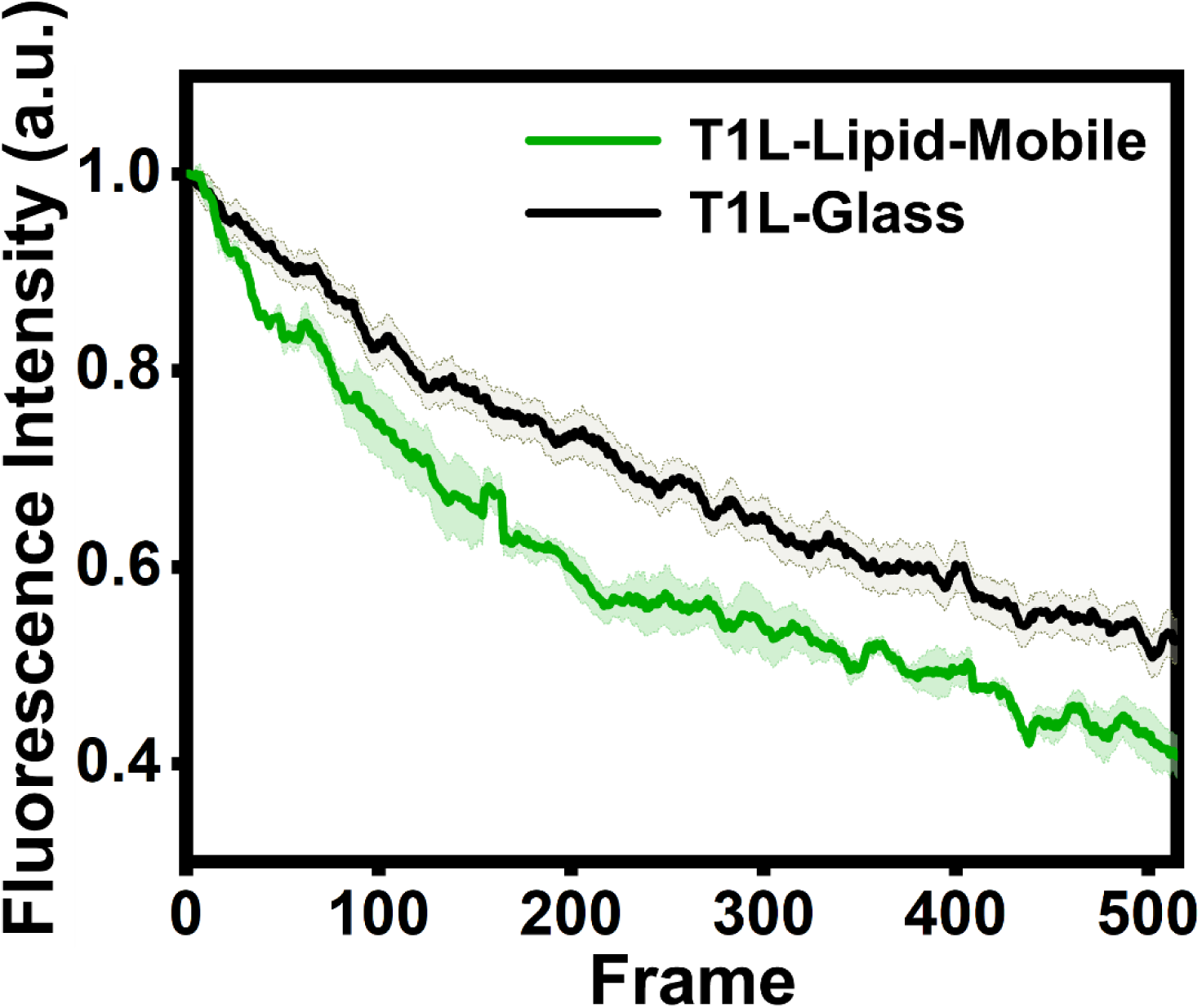
Capsid uncoating of T1L ISVPs during the interaction with lipids. Averaged line plots showing the fluorescence intensity of T1L ISVP on different surfaces over time. The error bar represents the standard error of the mean (S.E.M.).

**Figure S3.**
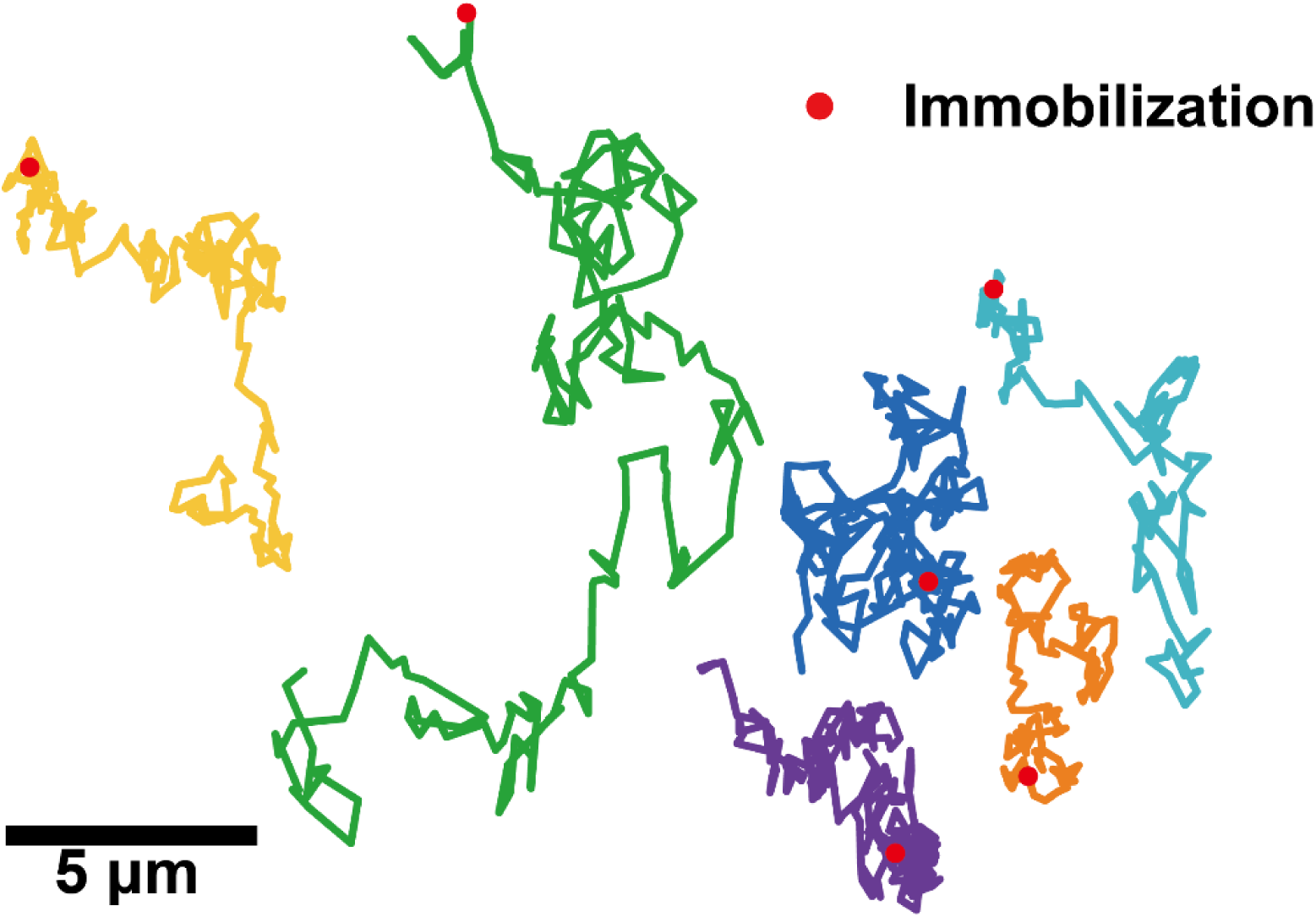
Complete immobilization of T1L/T3DM2 during imaging. Line plots showing the trajectories of T1L/T3DM2 that were completely immobilized on the membrane after interactions with lipids

**Figure S4.**
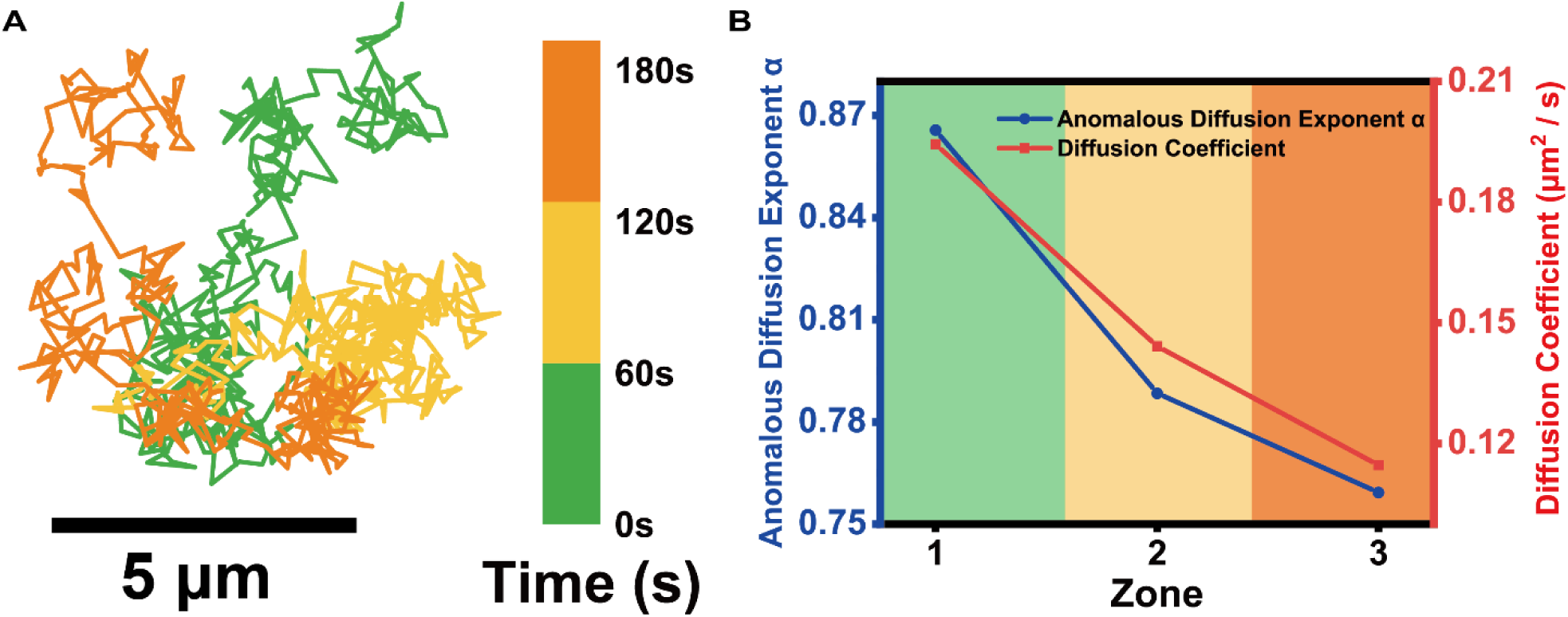
Confinement of T1L/T3DM2 ISVP increased over time on the planar-supported lipid bilayer. (A) Line plot showing a trajectory of a T1L/T3DM2 ISVP on the planar-supported lipid bilayer color-coded with time. (B) Line plots showing the corresponding anomalous diffusion exponent and diffusion coefficient change along with time from the trajectory shown in (A).

**Figure S5.**
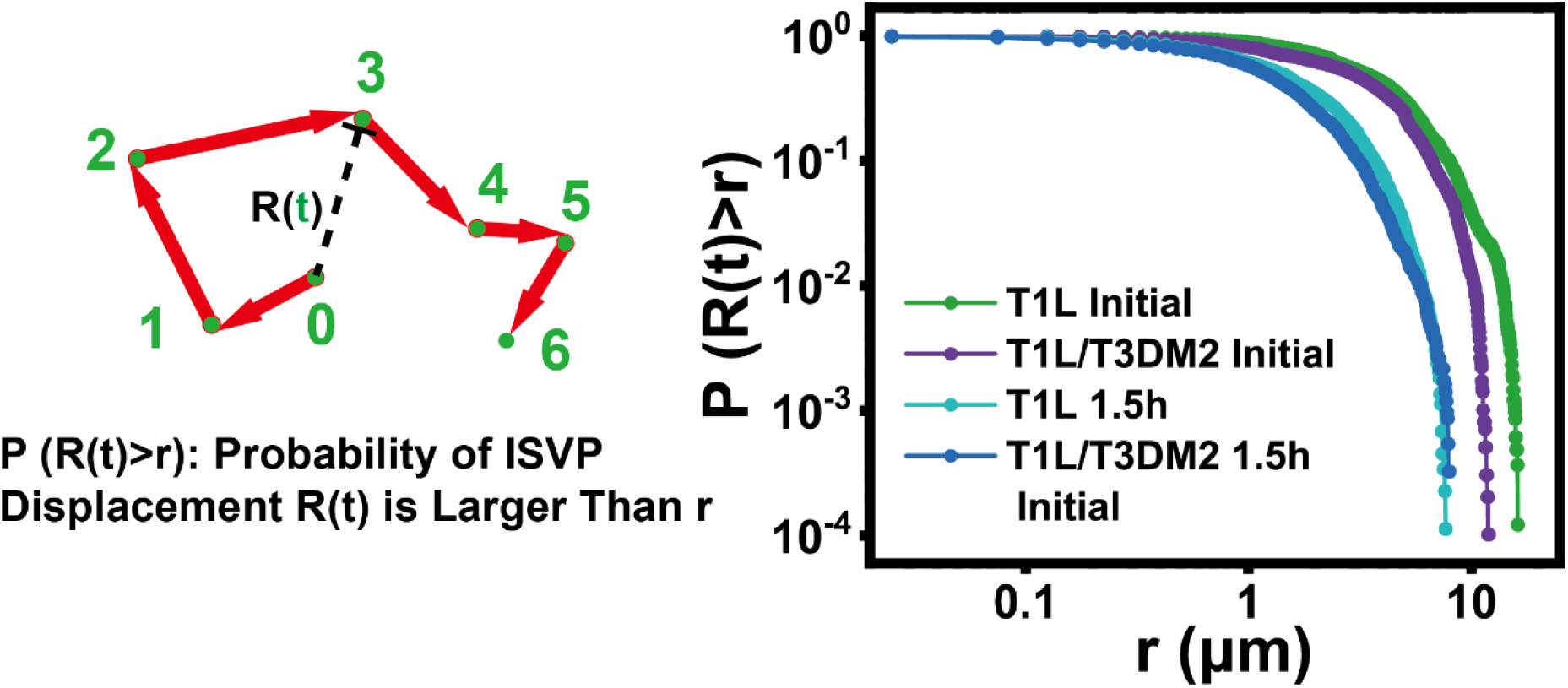
Probability of ISVP displacement. Schematic illustration of the displacement of ISVP R(t) in a trajectory (left) and log-log line plots (right) showing the probability of locating an ISVP outside of r distance away from the origin. T1L initial, n=8155; T1L/T3DM2 initial, n=9796; T1L after 1.5h, n=8786; T1L/T3DM2 after 1.5h, n=9196

**Figure. S6.**
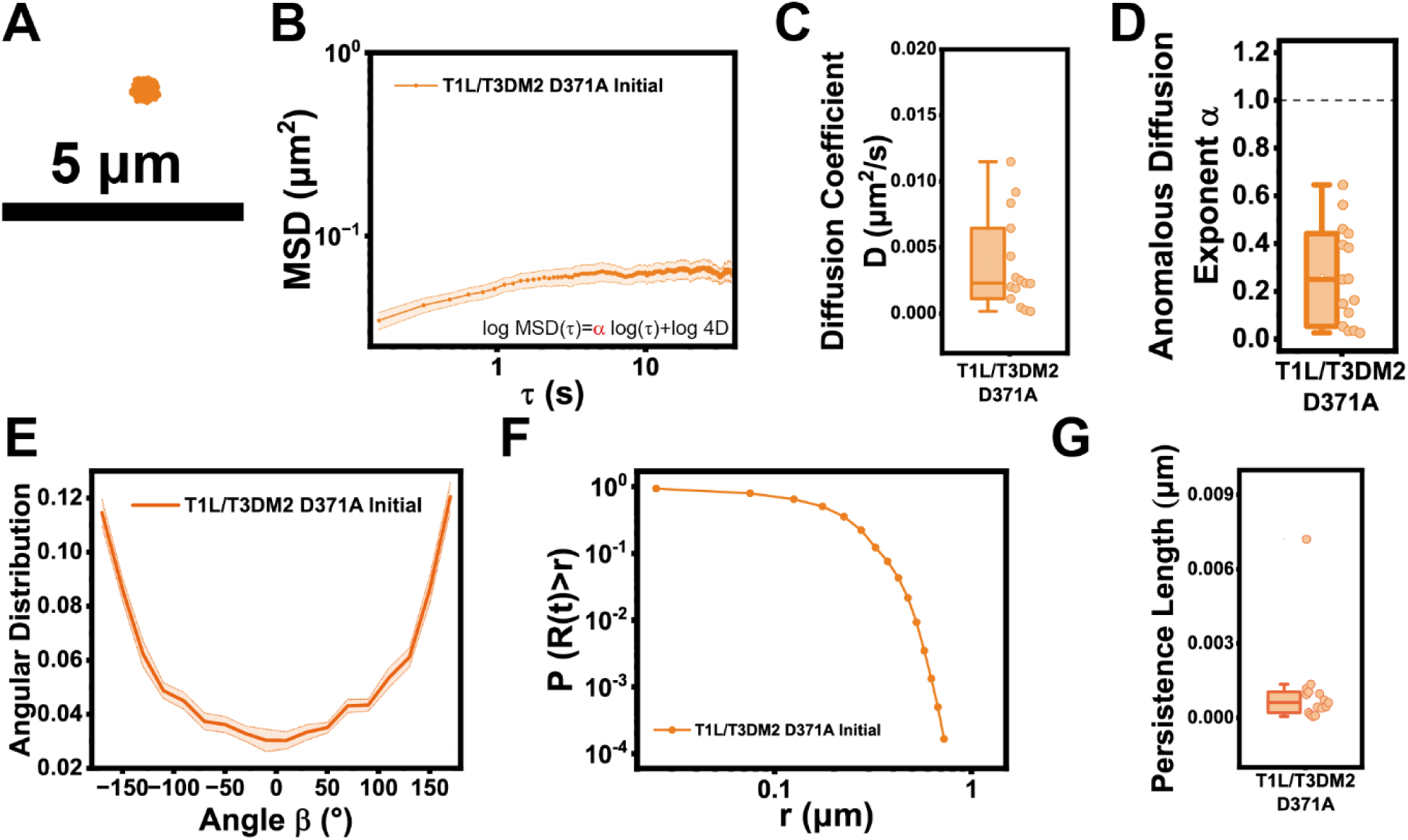
Translational diffusion of mutant ISVPs on the planar-supported lipid bilayer. (A) Representative line plots showing the trajectories of T1L/T3DM2 D371A ISVPs on the supported lipid bilayer. Scale bar, 5 μm. (B) Averaged log-log line plots showing the mean square displacement of ISVP trajectories. T1L/T3DM2 D371A initial (n=15). Error bar represents the S.E.M. (C and D) Statistical analysis of diffusion coefficient (C) and anomalous diffusion exponent (D) of trajectories from T1L/T3DM2 D371A initial (n=15). (E) Averaged line plot showing the angular distribution of turning angle in trajectories of T1L/T3DM2 D371A (n=15) ISVPs right after the recruitment of ISVPs on the planar-supported lipid bilayer. Error bar represents the standard error of the mean (S.E.M). (F) log-log line plots showing the probability of locating an ISVP outside of r distance away from the origin. T1L/T3DM2 D371A initial, n=5995. (G) Box and scatter plot of persistence length of trajectories from T1L/T3DM2 D371A initial (n=15). Each boxplot indicates the interquartile range from 25% to 75% of the corresponding data set. The mean and median are demonstrated as the square and the horizontal line, respectively. Statistical significance is highlighted by p values.

**Figure S7.**
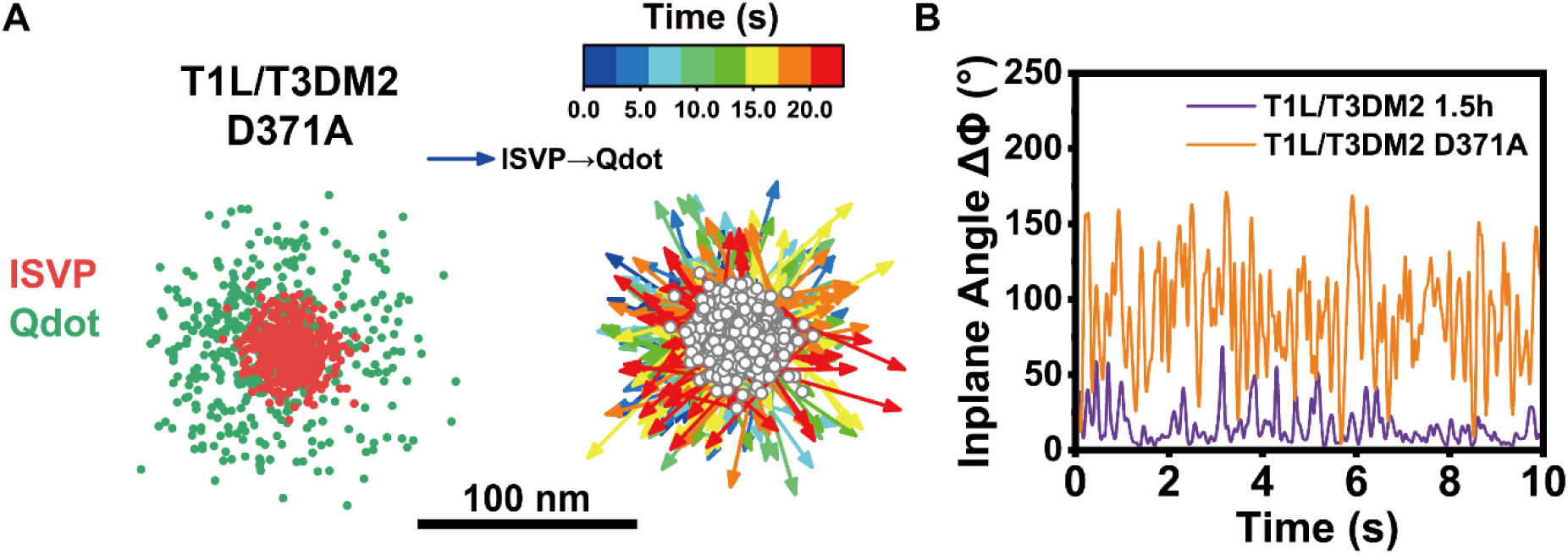
Rotational tracking of T1L/T3DM2 D371A ISVPs on the planar-supported lipid bilayer. (A) Representative rotational trajectories of T1L/T3DM2 D371A ISVP on the planar-supported lipid bilayer. Colormap encodes temporal information. (B) Line plots showing the change of in-plane angle in the trajectories shown in (A) and Fig. 5C.

**Figure. S8.**
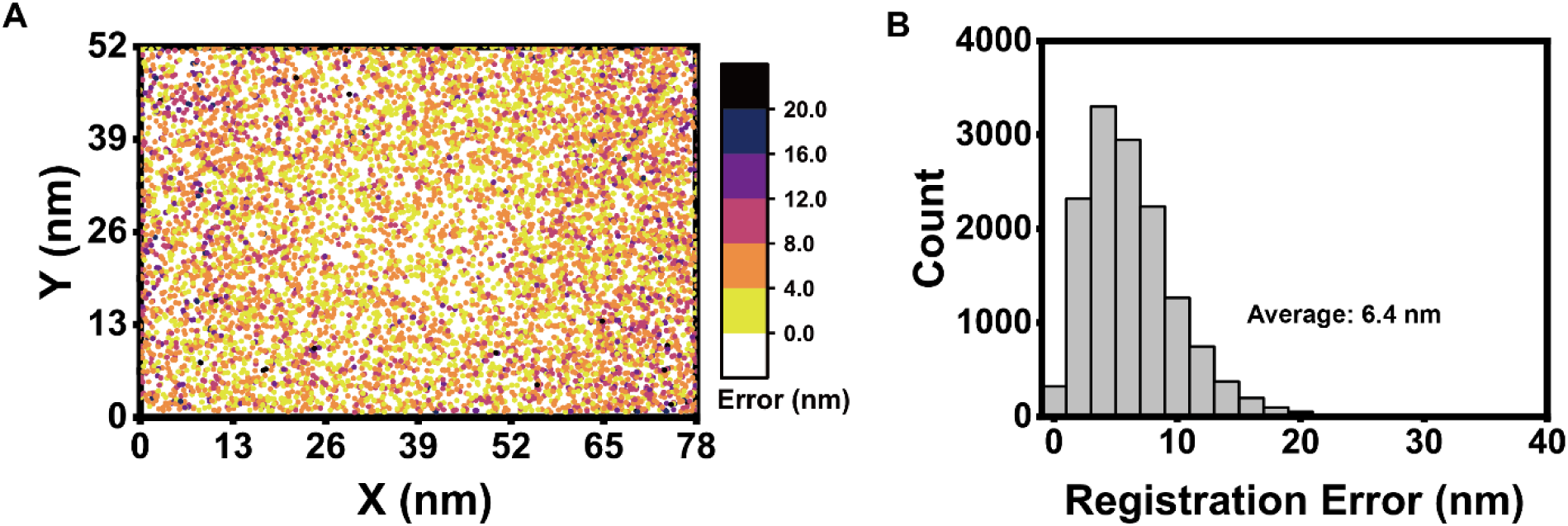
Registration performance of rotational Tracking. (A) A local registration error map showing the registration error of a set of fiducial markers after the local weighted color mapping. Colormap encodes the magnitude of the local mapping error at each location. (B) Histogram showing registration errors after a local weighted mapping was applied to a set of fiducial markers.

