## Supplemental Figures 1-8 for "Viral Capsid-Membrane Interactions Propel Non-Brownian Movements of Non-enveloped Reoviruses during Entry"

**Contents**

Supplementary figures S1-S8

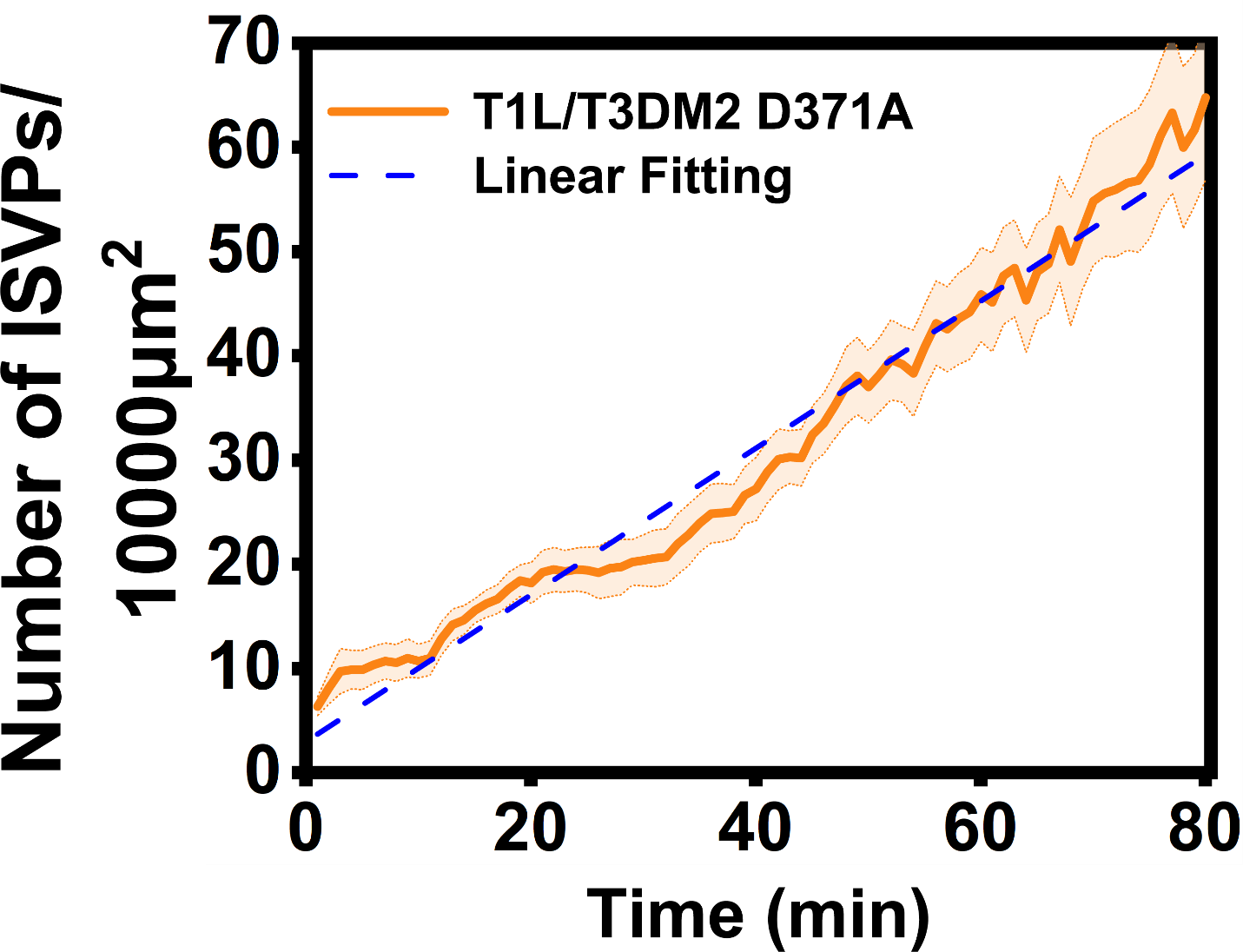

**Figure S1.** Recruitment of mutant reovirus ISVPs on the planar-supported lipid bilayer.

Averaged line plots showing the adsorption of 10 pM T1L/T3DM2 D371A ISVPs on supported lipid bilayer along time. Error bar represents the standard error of the mean (S.E.M).

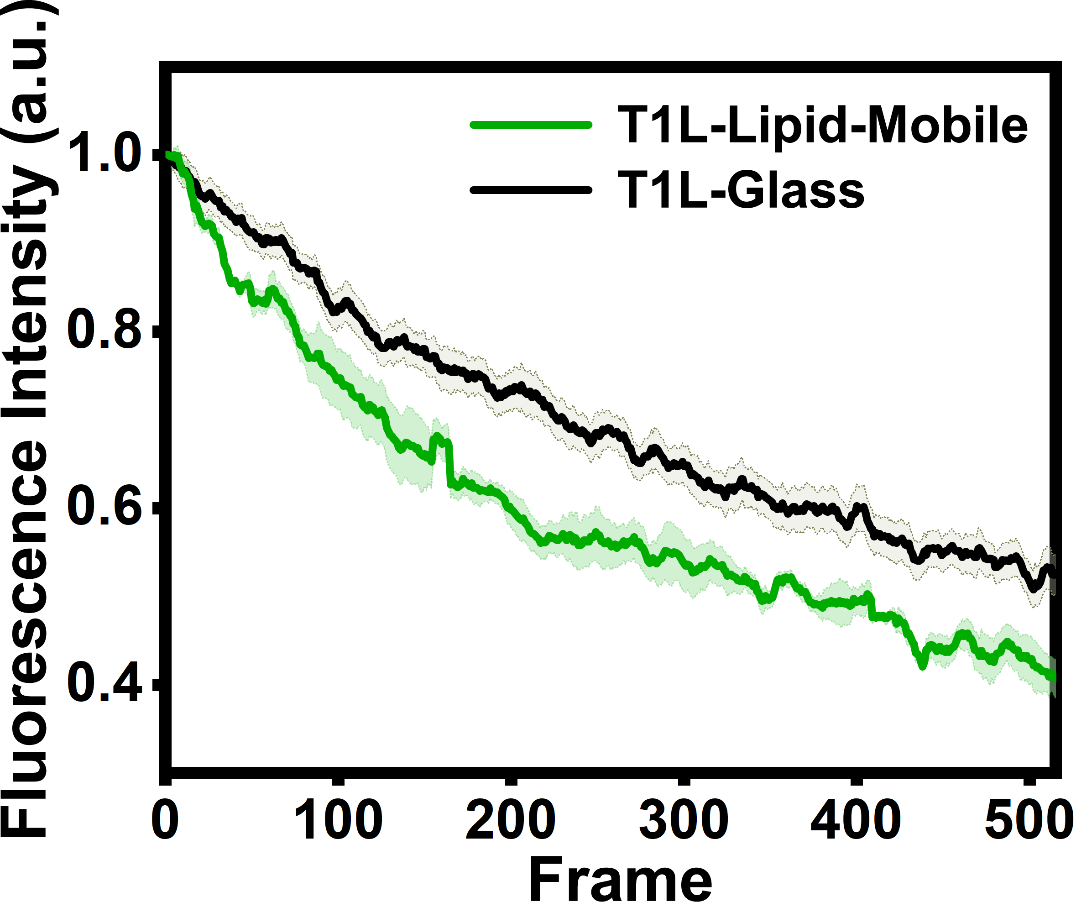

**Figure S2**. Capsid uncoating of T1L ISVPs during the interaction with lipids. Averaged line plots showing the fluorescence intensity of T1L ISVP on different surfaces over time. The error bar represents the standard error of the mean (S.E.M.).

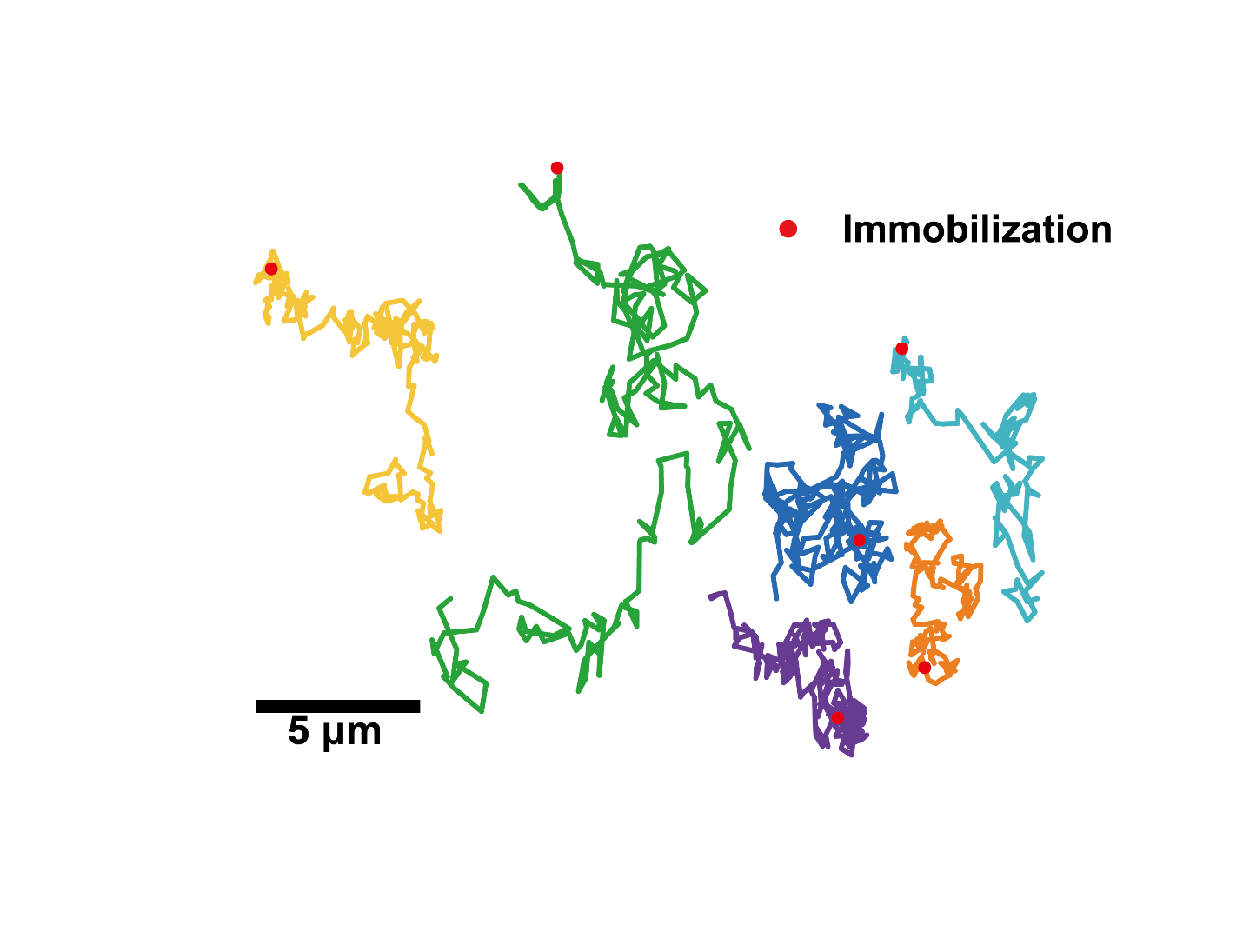

**Figure S3**. Complete immobilization of T1L/T3DM2 during imaging. Line plots showing the trajectories of T1L/T3DM2 that were completely immobilized on the membrane after interactions with lipids

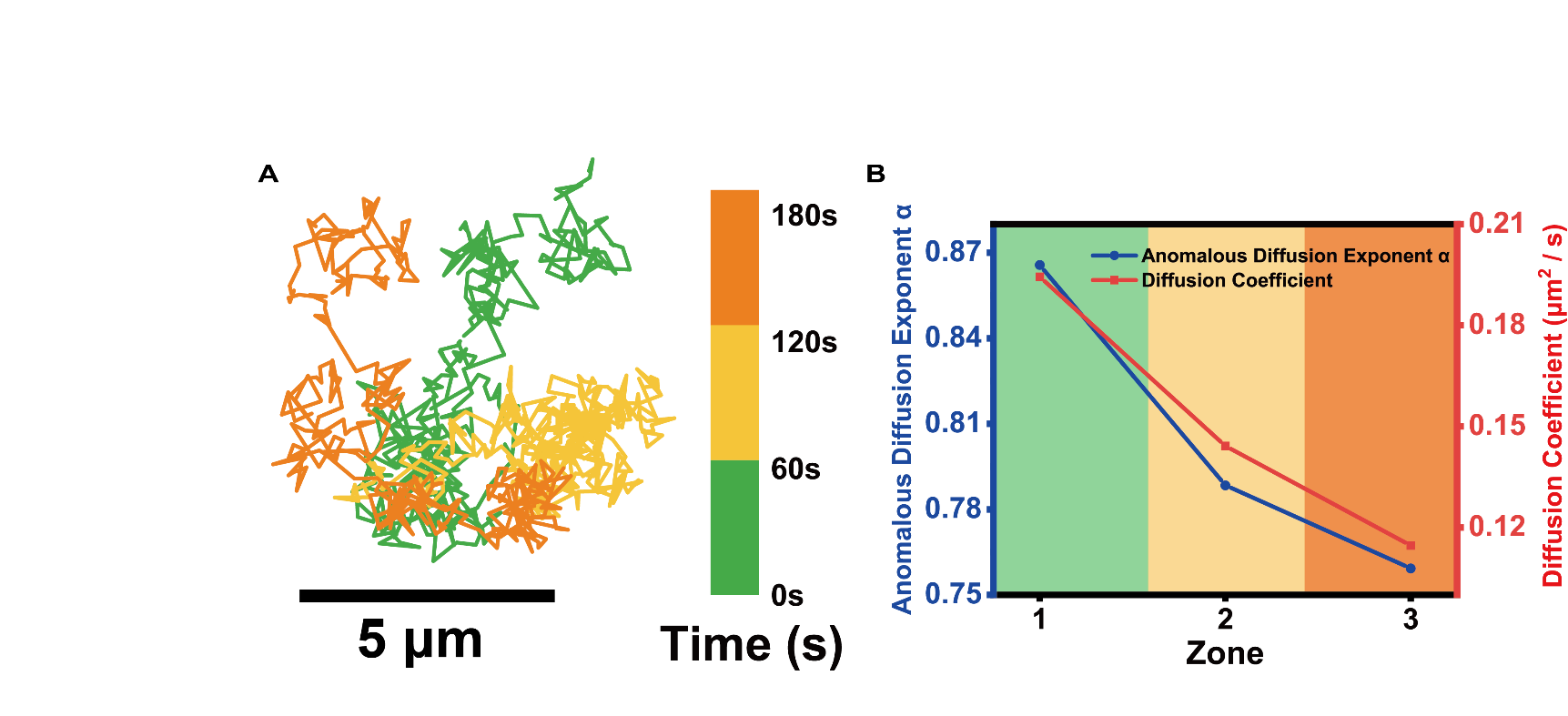

**Figure S4**. Confinement of T1L/T3DM2 ISVP increased over time on the planar-supported lipid bilayer. (A) Line plot showing a trajectory of a T1L/T3DM2 ISVP on the planar-supported lipid bilayer color-coded with time. (B) Line plots showing the corresponding anomalous diffusion exponent and diffusion coefficient change along with time from the trajectory shown in (A).

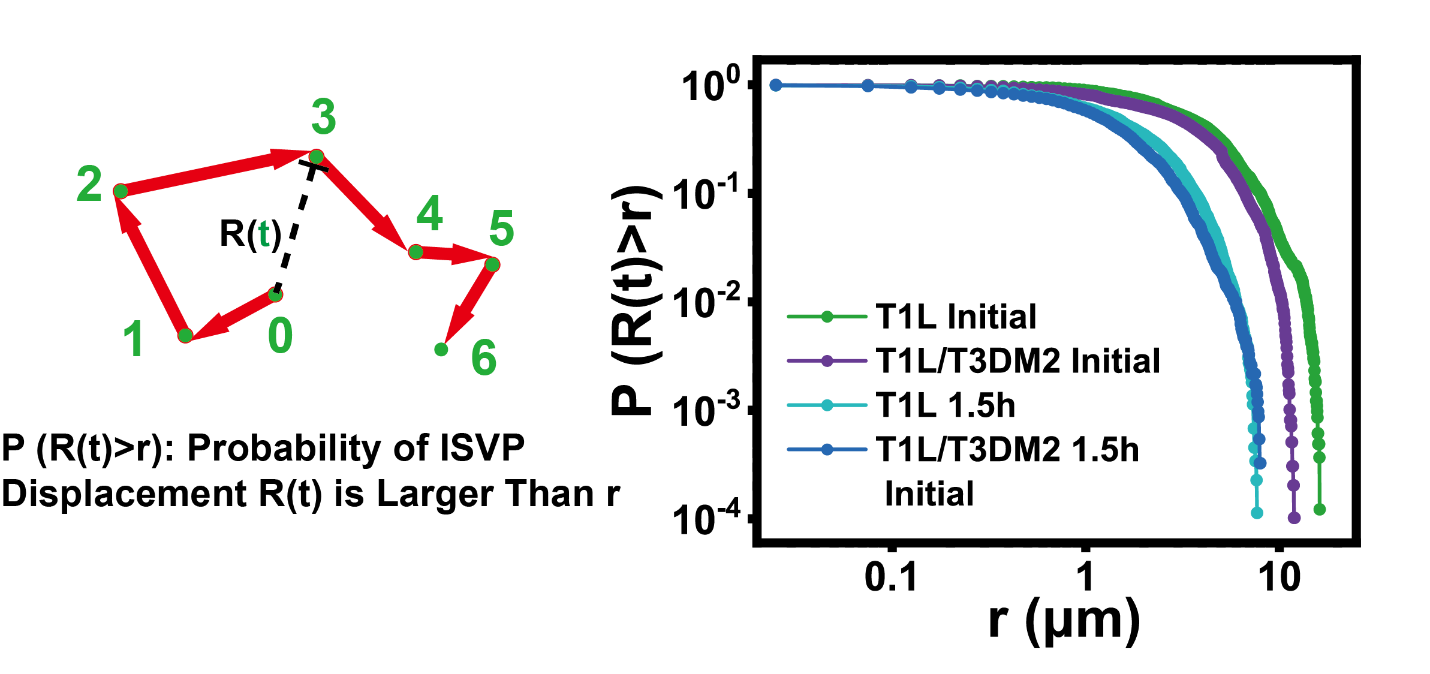

Figure S5. Probability of ISVP displacement. Schematic illustration of the displacement of ISVP R(t) in a trajectory (left) and log-log line plots (right) showing the probability of locating an ISVP outside of r distance away from the origin. T1L initial, n=8155; T1L/T3DM2 initial, n=9796; T1L after 1.5h, n=8786; T1L/T3DM2 after 1.5h, n=9196

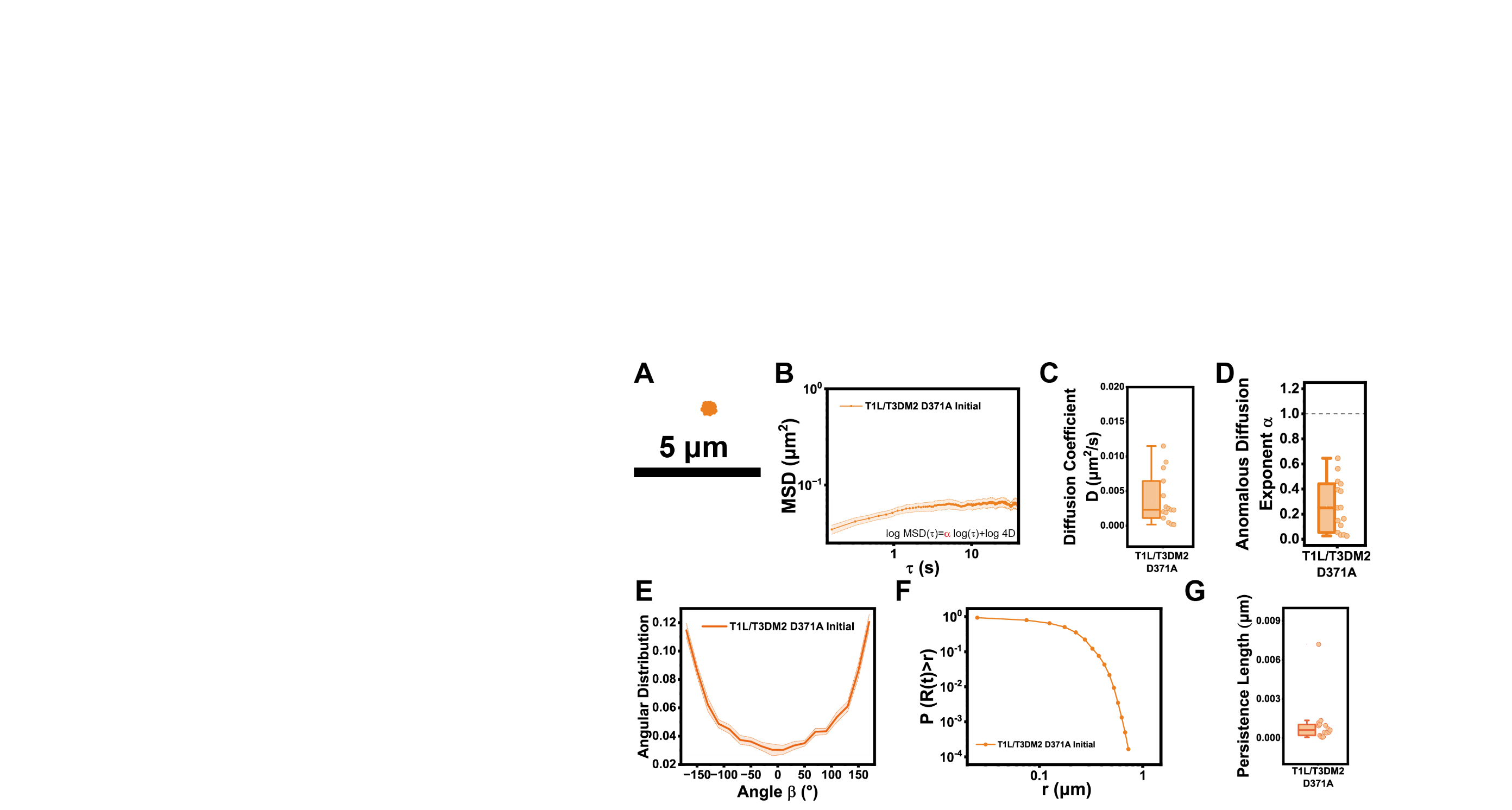

Figure. S6	Translational diffusion of mutant ISVPs on the planar-supported lipid bilayer.

(A) Representative line plots showing the trajectories of T1L/T3DM2 D371A ISVPs on the supported lipid bilayer. Scale bar, 5 μm.

(B) Averaged log-log line plots showing the mean square displacement of ISVP trajectories. T1L/T3DM2 D371A initial (n=15). Error bar represents the S.E.M.

(C and D) Statistical analysis of diffusion coefficient (C) and anomalous diffusion exponent (D) of trajectories from T1L/T3DM2 D371A initial (n=15).

(E) Averaged line plot showing the angular distribution of turning angle in trajectories of T1L/T3DM2 D371A (n=15) ISVPs right after the recruitment of ISVPs on the planar-supported lipid bilayer. Error bar represents the standard error of the mean (S.E.M).

(F) log-log line plots showing the probability of locating an ISVP outside of r distance away from the origin. T1L/T3DM2 D371A initial, n=5995.

(G) Box and scatter plot of persistence length of trajectories from T1L/T3DM2 D371A initial (n=15).

Each boxplot indicates the interquartile range from 25% to 75% of the corresponding data set. The mean and median are demonstrated as the square and the horizontal line, respectively. Statistical significance is highlighted by p values.

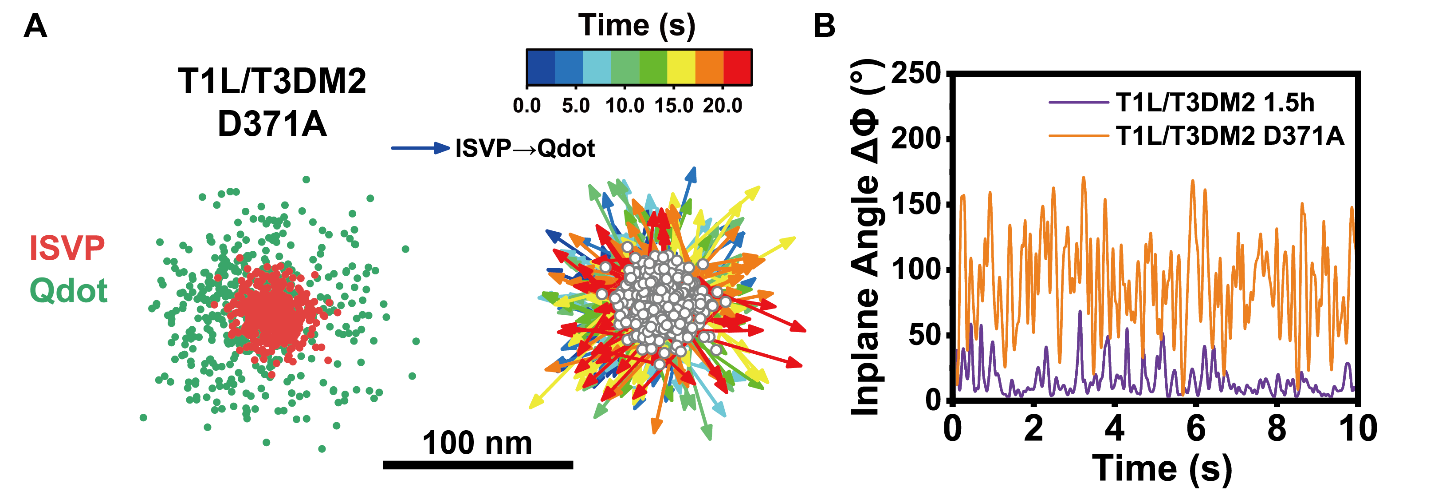

**Figure S7**. Rotational tracking of T1L/T3DM2 D371A ISVPs on the planar-supported lipid bilayer.

(A) Representative rotational trajectories of T1L/T3DM2 D371A ISVP on the planar-supported lipid bilayer. Colormap encodes temporal information.

(B) Line plots showing the change of in-plane angle in the trajectories shown in (A) and Fig. 5C.

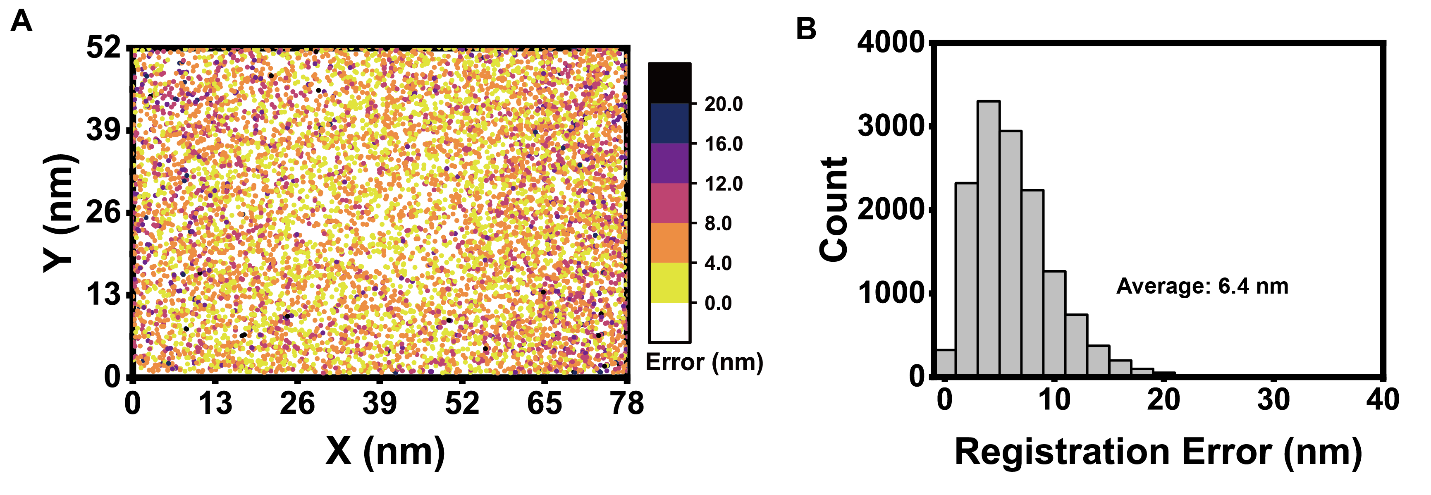

Figure. S8 Registration performance of rotational Tracking.

(A) A local registration error map showing the registration error of a set of fiducial markers after the local weighted color mapping. Colormap encodes the magnitude of the local mapping error at each location.

(B) Histogram showing registration errors after a local weighted mapping was applied to a set of fiducial markers.
